# Oxytocin Gα_i_ signaling-induced amygdala astrocytes processes retraction shapes behavioral stress response

**DOI:** 10.1101/2024.10.07.617014

**Authors:** Angel Baudon, Valentin Grelot, Ferdinand Althammer, Kai-Yi Wang, Clémence Denis, Syed Azmal Ali, Yudong Yan, Fernando Castillo-Diaz, Francesca Piacentini, Etienne Clauss Creusot, Volodya Hovhannisyan, Tim Schubert, Annabel C. Kleinwaechter, Jemima Helen, Tom Lakomy, Quirin Krabichler, Rachel Breton, Pierre-Alexis Derrien, Virginie Andry, Maria-Angeles Carrillo-de Sauvage, Valérie Demais, Aurélia Ces, Mélanie Kremer, Barbara Di-Benedetto, Yannick Goumon, Christian P. Schaaf, Valery Grinevich, Nathalie Rouach, Pascal Darbon, Inga D. Neumann, Antoine Adamantidis, Marta Busnelli, Lucile Benhaim, Jeroen Krijgsveld, Frank W. Pfrieger, Alexandre Charlet

## Abstract

Anticipated reactions to stressful situations are vital for the survival and well-being of organisms, and abnormal reactions are involved in stress-related disorders. The neuropeptide oxytocin is a key modulator ensuring well-adapted stress responses. Oxytocin acts on both neurons and astrocytes, but the molecular and cellular mechanisms mediating stress response remain poorly understood. Here, we focus on the amygdala, a crucial hub that integrates and processes sensory information through oxytocin-dependent mechanisms. Using an acute stress paradigm in mice, genetic and pharmacological manipulations combined with proteomic, morphological, electrophysiological and behavioral approaches, we reveal that oxytocinergic modulation of the freezing response to stress is mediated by transient Gαi-dependent retraction of astrocytic processes, followed by enhanced neuronal sensitivity to extracellular potassium in the amygdala. Our findings elucidate a pivotal role for astrocytes morphology-dependent modulation of brain circuits that is required for proper anticipated behavioral response to stressful situations.

## Introduction

Anticipated reactions to potentially dangerous situations are essential for animal survival, but their dysregulation can lead to pathological conditions such as post-traumatic stress disorders and chronic anxiety^1,2^. Rodents display similar responses to stress as stereotyped freezing behavior^3^. Understanding the cellular mechanisms mediating stressor detection and the resulting behavioral and physiological adaptations is crucial to devise effective treatments for anxiety and other stress-related disorders. Despite notable progress in this field, the cellular and molecular mechanisms mediating adequate physiological and behavioral responses to stress remain elusive.

Importantly, stress response requires coordinated activity of various brain regions. Among these, the amygdala plays a central role, as lesions of this brain structure impair sensory information interpretation across mammalian species, including rodents, monkeys, and humans^4^. Projections from the amygdala basolateral complex (BLA) to the central nucleus (CeA) are particularly critical for stress-related behavior^5^. Sensory information is integrated in the BLA and transmitted to the CeA which in turn modulates stress reactions as freezing behavior in rodents^3,6^. The activity of CeA neurons, especially in the laterocapsular part (CeL/C), is regulated by endogenous neuromodulators, notably oxytocin (OT)^7,8^.

OT is synthetized by neurons located in the hypothalamic accessory, supraoptic and paraventricular nuclei (PVN) that project to the pituitary and via axonal collaterals to various regions of the mammalian central nervous system^9^. OT modifies the balance between excitation and inhibition and regulates the salience of emotionally and socially relevant stimuli. In the context of the behavioral response to stress, this allows rapid switching between states of low and high arousal^10–12^. Within the CeL/C, OT modulates neuronal network activity and influences freezing. This highlights its role in regulating stress responses, and especially the acute stress-induced freezing in rodents^6,7,11–16^.

Remarkably, OT signaling involves not only neurons but also astrocytes as they express oxytocin receptors (OTR)^17^. Astrocytes are key regulators of neuronal function in the central nervous system^18,19^. They can detect and respond to neuronal activity by eliciting calcium transients; they release gliotransmitters, which in turn modulate synaptic transmission and they regulate neuronal excitability through neurotransmitter uptake and K^+^ buffering^20^. Consequently, astrocytes regulate the activity of neuronal circuits underlying behaviors in a brain-region specific manner^17,20^. Since OT is released in response to stressful conditions within the amygdala of rats^21^, this study aimed at determining whether and how astrocytes play a role in the oxytocinergic modulation of behavioral response to stress *in vivo*.

Here, we show that freezing response to acute stress is mediated by OTR- and Ca^2+^-dependent structural and functional changes in astrocytes, which in turn modulate neuronal activity in the CeL/C. Therefore, neuromodulatory signaling in astrocytes contributes to stress-induced neuronal hyperexcitability in the CeL/C.

## Results

To investigate the role of astrocytes in oxytocinergic modulation of behavioral response to stress *in vivo*, we adapted a paradigm of acute stress-induced freezing in adult mice^22^. Animals were exposed to five electrical foot shocks randomly delivered over a 10 minute period and freezing was quantified during the 20s preceding the shocks announced by an auditory cue (**Fig. 1a**).

**Figure 1:**
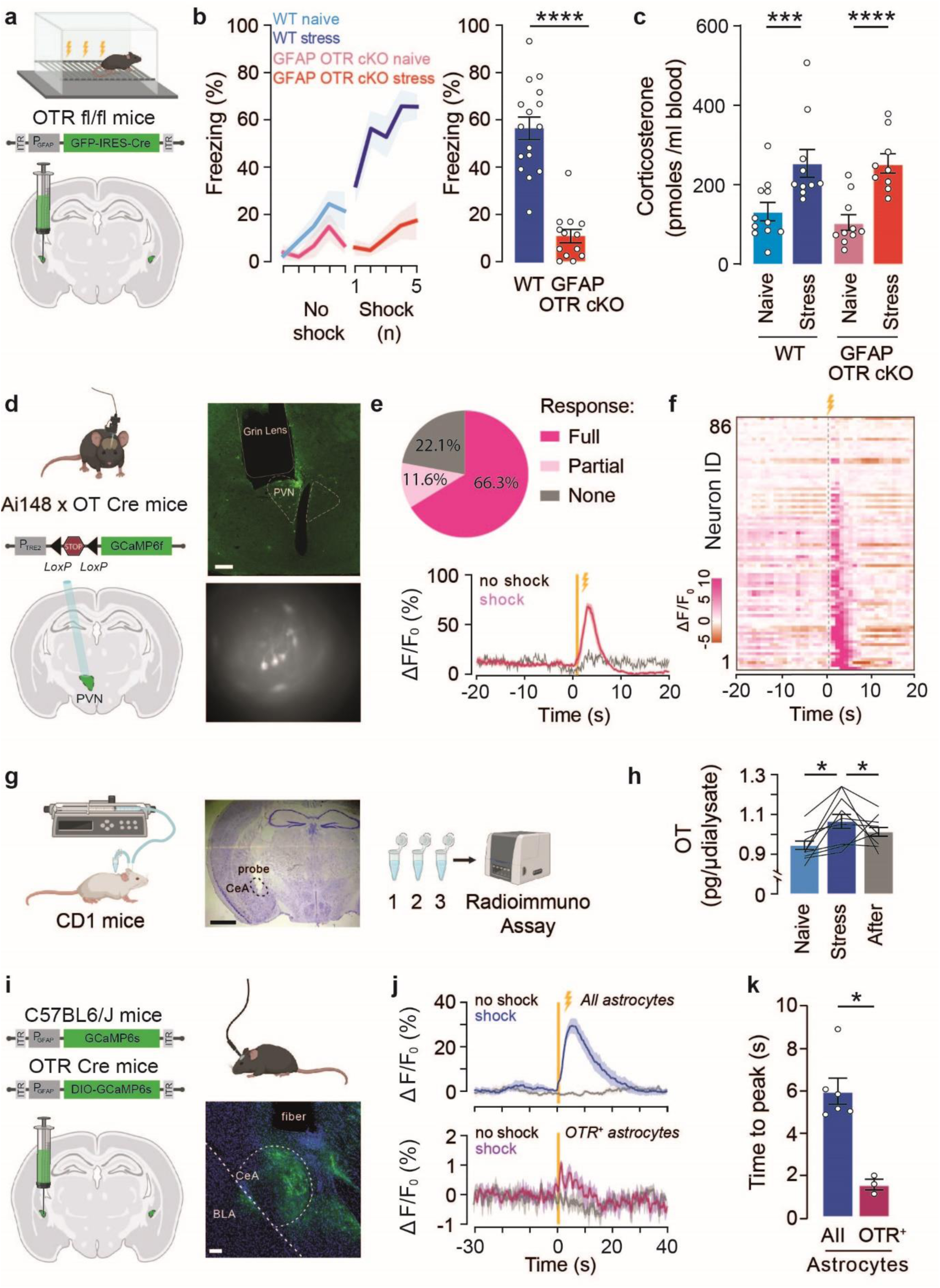
Impact of OTR deletion from CeL/C astrocyte on freezing behavior. **a.** Protocol for stress induction: Mice were submitted to five electrical foot shocks (1s, 0.6mA) at variable time intervals during a 10min session. Naïve mice were placed in the same cage without foot shocks. GFAP OTR cKO model: rAAV-GFAP-Cre was injected bilaterally in OTR-lox mice to knock out the OTR in astrocytes 3 weeks before experiments. **b.** Freezing behavior in WT and GFAP OTR cKO mice during the stress protocol. Left: Development of the freezing response during the habituation (Hab.) and after each electric shock exposure (Shock (n)). Right: quantification of the time spent freezing during the 20s following electric foot shock exposure in WT and GFAP OTR cKO mice. n_WT_=9 mice, n_GFAP OTR cKO_=11 mice. **c.** Plasma corticosterone levels in control and GFAP OTR cKO mice in naïve and stress male mice. n_WT_=11 mice, n_WT stress_=11 mice, n_GFAP OTR cKO_=10 mice, n_GFAP OTR cKO stress_=10 mice. **d.** Monitoring calcium activity in PVN OT neurons in freely moving animals. GCaMP6f was targeted to PVN OT neurons by crossing Ai148 mice with OT-Cre mice. GRIN lenses were implanted above the PVN and GCaMP fluorescence were captured thanks to a miniscope. Scale bar=100µm. **e.** Proportion of neurons showing calcium transients in response to each electric foot shock (full responders), to some electric foot shocks (partial responders) or showing no response (non responders). Peristimulus graph showing the mean variation of GCaMP6f fluorescence in PVN OT neurons before and after an electric foot shock. **f.** Scatter plot showing calcium responses to electric foot shocks of all PVN OT neurons tested. Calcium levels are encoded by color. n=4 mice. **g.** Microdialysis probe implanted in the amygdala allowed to obtain microdialysates in naïve condition, during the acute stress protocol and after mice return in the home cage. **h.** Quantification of the OT amout per microdialysate. **i.** Monitoring calcium activity of CeA astrocytes in freely moving mice. GCaMP was targeted to CeA astrocytes or to OTR-expressing CeA astrocytes following injection of rAAV-GFAP-GCaMP6s in WT mice (n=5) or rAAV-GFAP-DIO-GCaMP6s in OTR-Cre mice (n=3), respectively. Scale bar=100µm. **j.** Peristimulus graph showing the mean variation of GCaMP fluorescence in all CeA astrocytes (upper panel) or OTR-expressing astrocytes (lower panel). **k.** Bar plot showing mean time between the onset of electrical foot shock and the peak of astrocyte calcium response. Data are expressed as mean across animal ± SEM. Detailed statistics can be found in *Statistic Table 1*. * p<0.05, ** p<0.01, *** p<0,001, **** p<0.0001.

### Astrocyte OTR in the CeL/C contributes to stress-induced freezing

To directly assess whether astrocytic OT signaling influences the stress-induced freezing, we induced cell-specific loss of OTR by injecting an AAV encoding the Cre recombinase under the control of the glial fibrillary acidic protein (GFAP) promoter (rAAV-GFAP-Cre) to target astrocytes in the CeL/C of OTR fl/fl mice (GFAP OTR cKO, **Fig. 1a**)^17,23^. We observed that GFAP OTR cKO mice failed to exhibit freezing behavior during the stress session as compared with WT control mice (p<0.0001, **Fig. 1b** and **Extended data Fig. 1a-c**). Intriguingly, this effect was not observed in female mice (p=0.4494, **Extended data Fig. 1a-b**) indicating a putative sex-specific role of astrocytic OTR in the behavioral response to stress, which prompted us to focus on males. The absence of freezing behavior was not due to an altered physiological response to stress. Indeed, corticosterone levels in plasma were increased in stressed versus naïve animals in both WT and GFAP OTR cKO mice (p=0.6773, **Fig. 1c** and **Extended data Fig. 1d**).

Importantly, we tested whether the absence of freezing of GFAP OTR cKO mice could stem from changes in their nociceptive sensitivity. A comprehensive analysis of nociception and pain sensitivity in these animals revealed no differences in thermal cold (p=0.8080), thermal heat (p=0.2969) and mechanical (p=0.8534) nociception, nor in spontaneous pain as tested with the conditioned place preference test compared to WT mice (p=0.5774, **Extended data Fig. 1e-f**). We next examined whether the absence of freezing could be explained by GFAP OTR cKO mice displaying depressive-like symptoms. This appears unlikely as GFAP OTR cKO and WT mice performed similarly in the sucrose preference test and in the direct social interaction test (**Extended data Fig. 1g-h**).

### Acute stress induces PVN OT neurons activation, CeA OT release and astrocytes recruitment

Based on the behavioral characterization, astrocytes are involved in oxytocinergic modulation of stress-induced freezing *in vivo*. To decipher the underlying cellular mechanisms, we tested whether the acute stress paradigm activated PVN OT neurons and CeA astrocytes *in vivo*. To this end, we first recorded the calcium activity of OT neurons in the PVN in freely moving mice during the acute stress paradigm (**Fig. 1d**). To monitor individual neuronal calcium responses, we genetically targeted the calcium sensor GCaMP6f to OT neurons by breeding floxed GCaMP6f transgenic mice (line Ai148D) with OT-Cre mice^24^. Gradient-Index (GRIN) lenses were chronically implanted above the PVN for imaging of calcium activity using micro-endoscope in freely-moving mice (**Fig. 1d**). Electric foot shocks induced calcium transients in 78% of the GCaMP6f-expressing OT neurons (n=67/86; **Fig. 1e-f**). Interestingly, PVN OT neurons also responded to stress induced by tail suspension (p= 0.011) but not by cage shaking (p=0.750, **Extended data Fig. 1i-j**). These findings established that the acute stress protocol reliably activated OT neurons in the hypothalamus.

Next, we investigated whether acute stress triggered OT release in the CeA, which is a well-known target region of PVN OT neurons^7^. To this end, we a microdialysis probe within the CeA region to measure the release of endogenous OT in freely moving mice (**Fig. 1g**). We found that foot shock-induced stress induces an increase in CeA OT concentration (p=0.0162, **Fig. 1h**). This result showed that the foot shock-induced stress paradigm induces endogenous OT release in the CeA.

We then tested whether this acute stress paradigm induced a Ca^2+^ response in CeA astrocytes. To this end, we injected a rAAV carrying GCaMP6s under the control of the GFAP promoter (rAAV-GFAP- GCaMP6s) in the CeA and implanted optic fibers above the CeA to image Ca^2+^ activity in astrocytes population using fiber photometry in freely-moving mice (**Fig. 1i**). Similar to PVN OT neurons, electric foot shocks reliably increased Ca^2+^ transients in GCaMP6s-expressing astrocytes (**Fig. 1j-k** and **Extended data Fig. 1k-l**). To identify the contribution of the ∼20% of OTR-expressing astrocytes^17^ to this response, we injected OTR-Cre mice^25^ with a Cre-dependent AAV targeting astrocytes and expressing GCaMP6s. We observed that, in response to electric foot shocks, OTR-expressing astrocytes displayed Ca^2+^ transients that were approximately three times faster than the pan-astrocyte population in the CeA (p=0.023, **Fig. 1j-k** and **Extended data Fig. 1k-l**). These results established that the acute stress paradigm sequentially activated OTR-expressing astrocytes and the general astrocyte population in the CeA.

### Acute stress induces changes in amygdala proteome

To gain insight into the molecular and cellular mechanisms induced by acute stress, we performed label-free quantitative mass spectrometry proteomics on amygdala samples (**Fig. 2a**). Principal component analysis clearly separated naïve from stressed mice (**Fig. 2b**), indicating an extensive stress-induced remodeling of the amygdala proteome. Two large clusters of proteins were either up- or down-regulated upon stress (**Fig. 2c** and **Extended data Fig. 2**). Further analysis revealed that up- and downregulated proteins are involved in axonal activity and morphological remodeling of astrocytes (**Extended data Fig. 2a**), indicating that stress modifies protein levels in CeA astrocytes. Indeed, profound expression changes were observed among 59 astrocyte proteins (**Extended data Fig. 2b**), showing, among others, an increase in GFAP cytoskeleton protein. Analysis of gene ontology (GO) terms and protein-protein interactions revealed multiple signaling components, pathways, transmitter receptors and ion channels, most of them related to synaptic activity (**Fig. 2d**). Gene set enrichment analysis (GSEA) indicated that differentially expressed proteins are involved in multiple metabolic and signaling processes, as well as in structural cellular components (**Extended data Fig. 2b-c**). Among them, we found a significant change in intermediate filament proteins (**Fig. 2e-f**). Collectively, these data revealed that acute stress induces changes in astrocytes proteome, notably upregulation of intermediate filaments.

**Figure 2:**
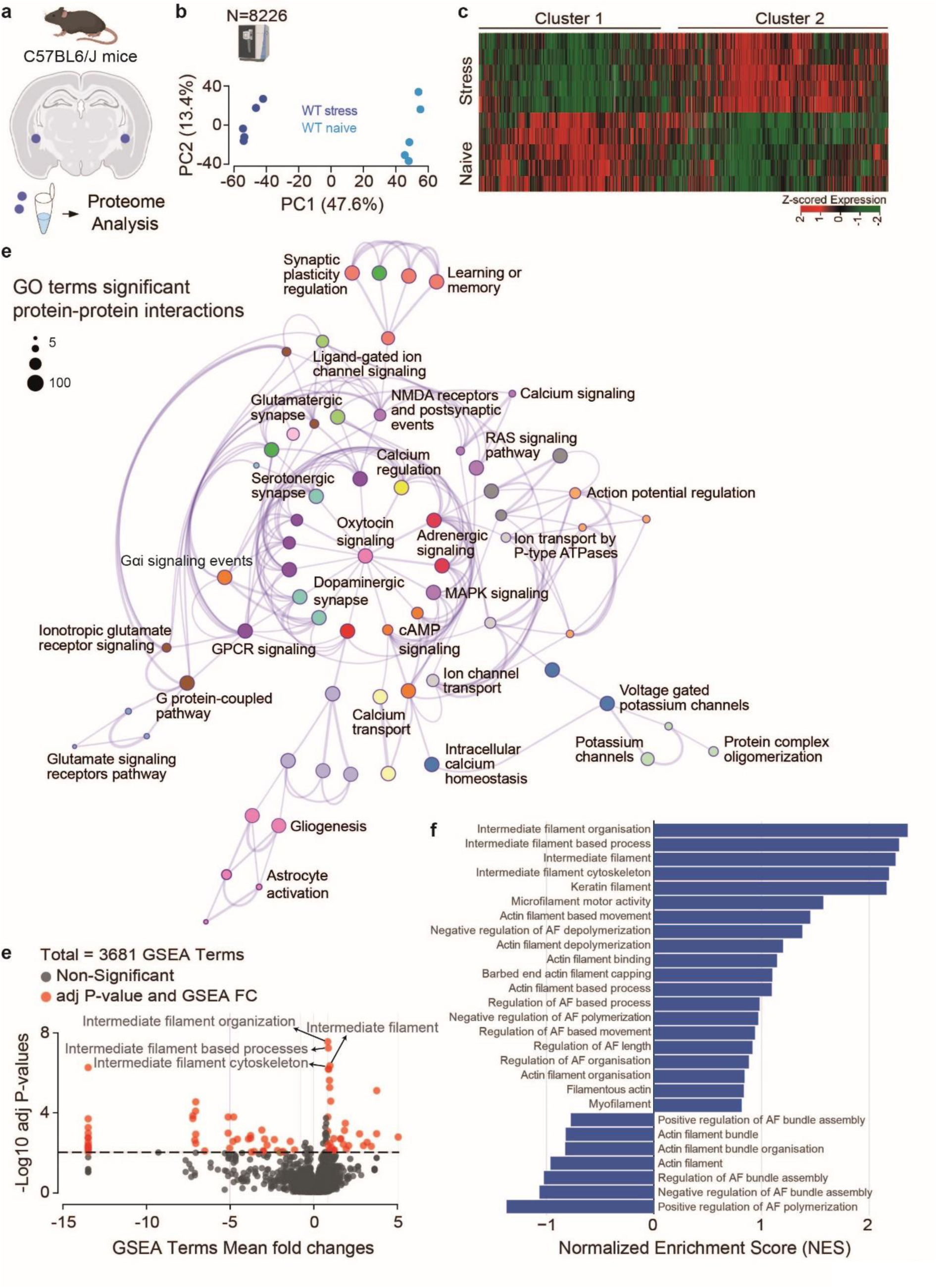
Proteome of the amygdala in naïve and stressed mice**. a.** Experimental design for proteome analysis. **b.** Principal component analysis (PCA) of the proteomic data. Dots represent proteomic data sets from individual control (sky blue) and stressed mice (dark blue). **c.** Heat map and hierarchical clustering of differentially expressed proteins. **d.** Network based on Protein-Protein Interaction (PPI) Map. Network of enriched terms obtained by analyzing top-300 genes by Metascape. Each node represents a term, and the node size is directly proportional to the number of input proteins grouped into each term. Node color denotes cluster identity. Gene ontology (GO) terms with a similarity score >0.5 are connected by an edge and the edge thickness represents the similarity score. **e.** 1D Enrichment Analysis of gene set enrichment analysis (GSEA) data. Each point represents a gene set obtained after GSEA analysis of the proteomic data, displayed as the average fold change of the proteins within that gene set as a function of the adjusted p-value of the GSEA analysis. Significant GSEA terms (adjusted p-value < 0.05) are highlighted in red. **f.** Bar plot of normalized enrichment scores (NES) obtained from GSEA analysis using the ranked fold changes and adjusted p-values for all the differential expressed proteins. The bar chart highlights the cytoskeletal-related pathways, with pathways exhibiting both positive and negative enrichment under stress conditions. Detailed statistics can be found in *Statistic Table 2*.

### OTR signaling mediates acute stress-induced changes in astrocyte morphology and K^+^-dependent neuronal depolarization

We next investigated whether the acute stress-induced changes in proteins entail changes in astrocyte morphology. To achieve this, we injected a rAAV-GFAP-GFP in the CeL/C of naïve and stressed mice. Using the IMARIS software, we performed three-dimensional reconstructions of GFP-expressing astrocytes from confocal images (**Fig 3a-b** and **Extended data Fig. 3a-b**). We found that the acute stress protocol decreases the morphological complexity of astrocytes, which displayed reduced process ramification as measured by Sholl analysis (p<0.0001, **Fig. 3b-c** and **Extended data Fig. 3c**). Notably, this morphological response was absent from GFAP OTR cKO mice (p=0.167, **Fig. 3b-c** and **Extended data Fig. 3c**), suggesting an OTR-dependent mechanism. Stress-induced changes in morphology were accompanied by an OTR-dependent increase in GFAP expression in CeL/C astrocytes (p<0.001, **Fig. 3d** and **Extended data Fig. 3d**). Consistent with this result, the GFAP volume measured by immunofluorescence was inversely correlated to astrocyte process ramification (r² = 0.569, p<0.0001, **Fig. 3e** and **Extended data Fig. 3e**). Together, these results revealed that acute stress concomitantly induces OTR-dependent upregulation of GFAP in astrocytes and a reduction of their morphologic complexity.

**Figure 3:**
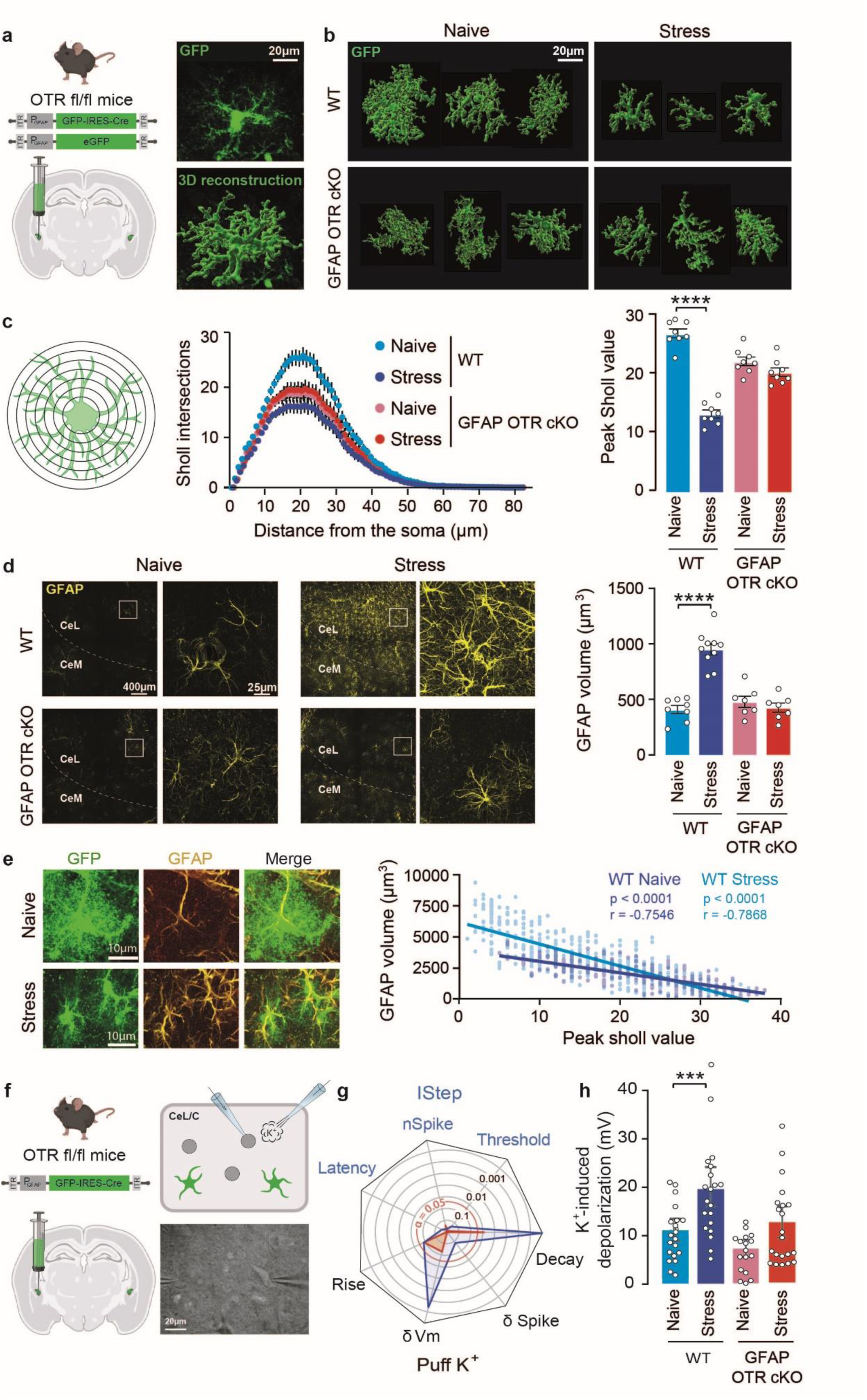
Morphological analysis of the CeL/C astrocytes network after stress. **a.** Imaging of CeL/C astrocytes in different mouse lines using targeted expression of GFP using rAAV-GFAP-GFP injection. Right: reconstruction of CeL/C astrocytes based on their GFP expression. Scale bar=20µm. **b.** Representative images of reconstructed astrocytes from WT and GFAP OTR cKO mice under naïve and stress conditions. Scale bar=20µm. **c.** Estimation of astrocyte ramification using Sholl analysis. Left: Diagram depicting the Sholl method. Middle: Sholl plot showing the number of astrocyte processes crossing the concentric Sholl circles in function of the distance from the soma. n=8 mice per group, n_naïve_=4032, n_stress_=3402, n_naïve GFAP OTR cKO_=2843, n_stress GFAP OTR cKO_=2968 astrocytes. Right: Comparison of astrocytes ramification in different experimental groups based on mean maximum Sholl values. **d.** Quantification of GFAP volume after immunohistochemical staining and 3D reconstruction. Scale bar=400 and 25µm. **e.** Correlation between astrocyte ramification revealed by GFP and their GFAP volume. n_naïve_=333 astrocytes, n_stress_=489 astrocytes. Scale=10µm. Scatter plot showing a negative correlation between GFAP volume and peak Sholl values in individual astrocytes. **f.** Astrocyte-specific cKO of OTR following rAAV-GFAP-Cre injection in the CeL/C of OTR-lox mice 3 weeks before stress experiments. Whole-cell patch-clamp recordings from CeL/C neurons in acute slices. Micrograph of slice during patch-clamp experiment. Scale bar=50µm. **g.** Spider plot showing the log p-values given by the comparison of different electrophysiological features. Current steps (IStep) were used to measure the spike-firing threshold, the number of spikes at the second active step (nSpike), and the latency to the first spike. K^+^ puff (30mM in aCSF), was used to measure the difference in spike firing (δSpike), the amplitude of the K^+^-induced depolarization (δVm), and its rise/decay constants. **h.** Amplitude of neuronal membrane depolarization during the K^+^ puff. n=18-21 neurons. In bar graphs, data are expressed as mean across animal ± SEM. Detailed statistics can be found in *Statistic Table 3*. * p<0.05, ** p<0.01, *** p<0.00

Given that astrocytic OTR are necessary for acute stress-induced freezing (**Fig. 1**) and knowing that astrocytic regulation of neuronal excitability can depend on their morphology^26,27^, we characterized the excitability of CeL/C neurons in brain slices of naive and stressed mice using whole-cell patch-clamp recordings (**Fig. 3f**). We found that current injections induced similar voltage responses in neurons from WT naïve and stressed mice (p_rheobase_=0.300, p_latency to spike_=0.372, p_number of spikes_=0.521, **Fig. 3g** and **Extended data Fig. 3f**). However, K^+^ puff (30mM)-induced depolarization of neuronal membrane potential (**Fig. 3g-h** and **Extended data Fig. 3g**) was higher in stressed mice compared to naïve mice (p=0.0002). The kinetics of this depolarization was modified at the level of decay (p_rise_=0.086, p_decay_<0.0001, **Fig. 3g** and **Extended data Fig. 3**) and the K^+^ puff did not elicit higher spike frequency in neurons from stressed versus naive animals (p_number of spikes_=0.094, **Fig. 3g** and **Extended data Fig. 3**). Importantly, we found that this stress-induced increase in K^+^-dependent neuronal depolarization was not significant in GFAP OTR cKO, as compared to naive mice (p_depolarization_=0.062, p_rise_=0.088, p_decay_=0.025, p_number of spikes_=0,676, **Fig. 3g-h** and **Extended data Fig. 3g**). Taken together, the results revealed that OTR-dependent signaling in astrocytes mediates acute stress-induced morphologic changes in astrocytes and an increase of K^+^-dependent depolarization of CeL/C neurons.

### Stress-induced changes in astroglial coverage of synapses and regulation of their efficacy within the amygdala CeL/C

The acute stress-induced changes in astrocyte morphology raised the question of whether this also affected the astrocytic coverage of synapses. This is of physiologic importance as such changes can influence circuit function^28^. To address this, we injected rAAV-GFAP-GFP in the CeL/C to label astrocytes. We then stained for vGluT1, vGluT2, and Homer1 to visualize pre- and postsynaptic elements of excitatory glutamatergic synapses (**Fig. 4a-b** and **Extended data Fig. 4a**). Confocal microscopy revealed a significantly reduced number of synapses contacted by astrocytes from mice subjected to acute stress compared to those from naïve mice (p=0.001, **Fig. 4c**). These results suggest that acute stress prompted the retraction of astrocytic processes away from synapses. Notably, this difference was not observed in GFAP OTR cKO mice, indicating that this change in synaptic coverage was mediated by astrocytic OTR (p=0.998, **Fig. 4c**). As a complementary method to visualize the coverage of synapses by astrocytic processes, we used transmission electron microscopy. In support of our observations by light microscopy, we observed a lower density of astrocytic processes surrounding synapses in CeL/C from stressed mice compared to naïve ones (p<0.0001; **Extended data Fig. 4b**). Together, these observations indicated that acute stress initiated an OTR-dependent retraction of astrocytic processes contacting synapses in the CeL/C.

**Figure 4:**
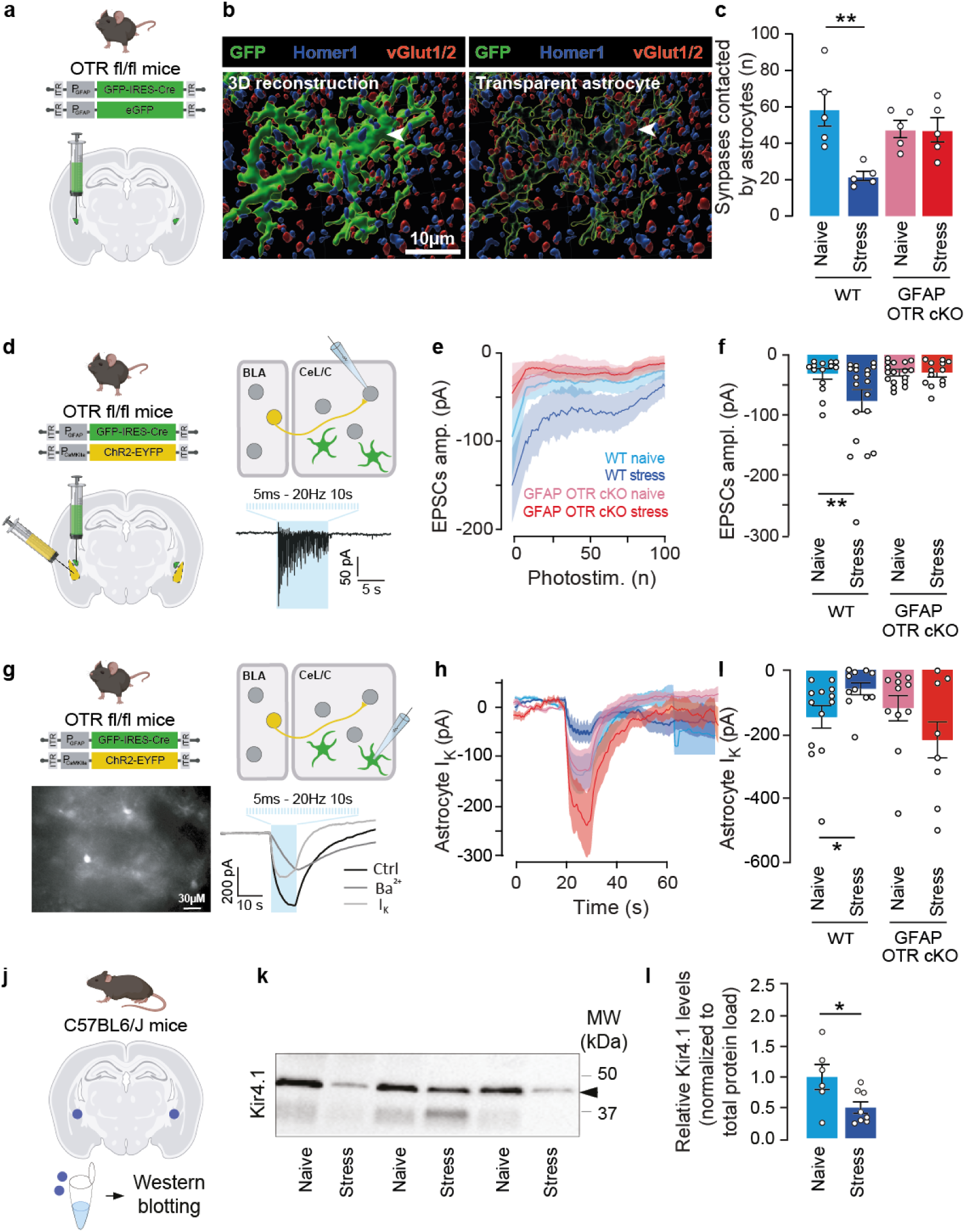
Involvement of astrocytic OTR on stress-induced modification of CeL/C synaptic coverage and BLA-to-CeL/C excitatory synaptic transmission. **a.** Imaging of CeL/C astrocytes using targeted expression of GFP using rAAV-GFAP-GFP. **b.** Representative micrograph showing synapses and surrounding astrocytic processes following IMARIS-based 3D reconstruction. **c.** Quantification of the contacts between synapses and astrocyte processes per reconstructed astrocyte. n=5 animals in each group. **d.** Analysis of BLA-to-CeL/C synaptic transmission using optogenetic stimulation of the BLA axons and recordings of the EPSCs in CeL/C neurons in naïve and stressed mice. AAV-CaMKII-ChR2 was injected in the BLA of mice and 3 weeks after, we measured the currents evoked by BLA neurons activation in CeL/C neurons of presynaptic BLA neurons. Representative recording of EPSCs in CeL/C neurons evoked during photostimulation. **e-f.** Time course and quantification of photostimulation-evoked EPSCs amplitude. n_WT naïve_=14, n_WT stress_=19, n_GFAP OTR cKO naïve_=17, n_GFAP OTR cKO stress_=13 neurons. **g.** Experimental approach to record K^+^ currents in CeL/C astrocytes following optogenetic stimulation of BLA neurons in naïve and stressed mice. AAV-CaMKII-ChR2 was injected in the BLA of mice and 3 weeks after, the currents evoked by BLA neuron activation were measured in CeL/C astrocytes. **h-i.** Time course and quantification of photostimulation-evoked astrocytic current induced by BLA neurons stimulation before and after a 15min incubation in Ba^2+^ (200µM). The subtracted K^+^ current is shown in green. n_WT naïve_=13, n_WT stress_=11, n_GFAP OTR cKO naïve_=11, n_GFAP OTR cKO stress_=8 astrocytes. **j.** Quantification of K_ir_4.1 levels in naïve and stressed mice using 1mm punches of mice CeA were done on 400µm slices from naïve or stressed mice. **k.** Image of representative Western blot reacted with antibody against K_ir_4.1 in six animals from indicated experimental groups. **l.** Quantification of K_ir_4.1 levels in amygdala punches. n_naïve_=6, n_stress_=8 mice.Data are expressed as mean across animal ± SEM for panel c and m, and as mean across cell ± SEM for panels c, f, I, and l. Detailed statistics can be found in *Statistic Table 4*. * p<0.05, ** p<0.01.

We next tested whether the reduced coverage of synapses by astrocytic processes induced by acute stress affected their transmission properties. Previous studies suggested a major role of basolateral amygdala (BLA) to CeL/C projections in the adaptation to stress-induced freezing^28,29^. To optogenetically activate BLA neurons, we injected rAAV-CaMKII-ChR2-EYFP in the BLA (**Fig. 4d**). After visual verification of the BLA location of EYFP expression, we stimulated ChR2-expressing BLA neurons by light (5ms pulse, 20hz during 5s) and recorded the synaptic response of CeL/C neurons using patch-clamp recordings in acute brain slices of both naïve and stressed animals (**Fig. 4d** and **Extended data Fig. 4c-g**). Acute stress increased the overall amplitude of the evoked excitatory postsynaptic currents (EPSCs) in CeL/C neurons as compared to naïve mice (p_amplitude_=0.037, **Fig. 4e-f**), similarly to^30^. To test the involvement of OTR signaling in astrocytes, we performed the same recordings in GFAP OTR cKO mice. Under these conditions, the stress-induced increase in synaptic responses was abolished (p=0.948, **Fig. 4e-f**). Interestingly, the amplitude of EPSCs triggered by the two first photostimulations was not different between naïve and stressed mice (**Fig. 4e**, p=0.107). However, the BLA-evoked EPSCs amplitudes in the middle of the stimulation train was increased in stressed compared to naïve mice (**Fig. 4f**, p=0.031), indicating an incremental mechanism. This could arise from an accumulation of extracellular K^+^ with repeated stimulations.

To test the implication of astrocytic K^+^ buffering in this phenomenon, we recorded K^+^ currents in CeL/C astrocytes following photostimulation of BLA neurons. We patch-clamped SR101-labelled astrocytes (**Fig. 4g**) and verified their astrocytic identity based on their electrophysiological response (IV curve, **Extended data Fig. 4h**)^31^. To isolate the current due to K^+^ buffering, we subtracted from the current evoked by photostimulation of BLA neurons a second current recorded in the presence of Ba^2+^ (200µM) to block K^+^ currents (**Fig. 4g**). This experiment revealed that the amplitudes of stimulation-induced K^+^ currents were significantly reduced in stressed mice compared with naïve mice (p=0.030, **Fig. 4h-i**), indicating impaired K^+^ buffering in astrocyes. Since we cannot rule out a potential Ba^2+^ effect on neurons, we reproduced the experiment in GFAP OTR cKO mice. Crucially, the stimulation-induced K^+^ currents were not altered in GFAP OTR stressed mice compared with naïve mice (p=0.159, **Fig. 4h-i**). A decrease in astrocyte K^+^ buffering capacity could arise from their process retraction but also from reduced expression of inwardly rectifying K_ir_4.1 channels^19,32^. Thus, we extracted membrane proteins of CeA samples from stressed and naïve mice and detected K_ir_4.1 protein levels by western blot analysis (**Fig. 4j** and **Extended data Fig. 4i-j**). We found that the acute stress paradigm reduced K_ir_4.1 levels in CeA samples as compared with naïve controls (**Fig. 4j-l**). These results suggested that acute stress induces a retraction of astrocytic processes from synapses and diminishes the K^+^ buffering capacity at the BLA-to-CeL/C synapses, thereby modifying the strength of this connection.

These findings suggest that acute stress initiates an OTR-dependent retraction of perisynaptic astrocytic processes, leading to a diminished K^+^ buffering capacity at the BLA-to-CeL/C synapses, enabling adequate behavioral stress response (**Fig. 4m**).

### Identification of G proteins mediating OTR-dependent effects in astrocytes

We next aimed to identify the signaling pathways downstream of OTR underlying changes in Ca^2+^ transients in astrocytes after acute stress.

Previous proteomic analysis suggested that acute stress modifies GPCR and cAMP signaling pathways (**Fig. 2d**, **S2c** and **Extended data Fig. 5a**). Thus, we studied the subtype of G-proteins activated by OTR in astrocytes using bioluminescence resonance energy transfer (BRET)-based biosensors^33^. We found that upon OT binding, OTR couple to both Gαi and Gαq in cultured astrocytes (**Extended data Fig. 5b-d**). We next identified the G proteins coupling OTR to Ca^2+^ transients in CeL/C astrocytes (**Fig. 5**) by combining OGB1-based calcium imaging and pharmacological manipulations. We applied biased OTR agonists that favor either Gαi (atosiban) or Gαq (carbetocin) pathways or a full agonist ([Thr^4^,Gly^7^]- oxytocin, TGOT) that activates both pathways^33,34^. We then measured calcium transients in SR101- positive astrocytes in brain slices in the presence of TTX to block action potential-dependent neuronal activity (**Fig. 5a**). All three agonists increased the frequency and AUC of Ca^2+^ transients compared to baseline activity (frequency: p_TGOT_=0.024, p_atosiban_=0.001, p_carbetocin_=0.003; AUC: p_TGOT_=0.002, p_atosiban_=0.0007, p_carbetocin_=0.0008, **Fig. 5b** and **Extended data Fig. 5b**). Similar proportions of cells responded to each agonists (∼50%, **Fig. 5c** and **Extended data Fig. 5e**). The Gαq protein blocker YM254890 (500nM, 30min pre-incubation) prevented the carbetocin- and TGOT-induced calcium signaling (Carbetocin: Ca^2+^ transient frequency: p=0.296, TGOT: Ca^2+^ transient frequency: p=0.561, **Extended data Fig. 5f**). The Gαi protein blocker pertussis toxin (PeTX, 5µg/mL, 5h pre-incubation) abolished calcium activity induced by atosiban (Ca^2+^ transient frequency: p=0.316, **Extended data Fig. 5g**), while the response to TGOT remained unaffected (Ca^2+^ transient frequency: p=0.006, **Extended data Fig. 5g**). In order to characterize the Ca^2+^ transients induced by OTR-Gαi and OTR-Gαq pathways, we compared the parameters of calcium event triggered by different G protein activation. We found that the rise constant and amplitude of Ca^2+^ transients induced by atosiban were lower as compared to those induced by carbetocin (p_rise_=0.016, p_amplitude_= 0.013, **Fig. 5c-d**). Taken together, these pharmacological data indicated that activation of Gαi and Gαq proteins triggers distinct Ca^2+^ transient signatures in astrocytes.

**Figure 5:**
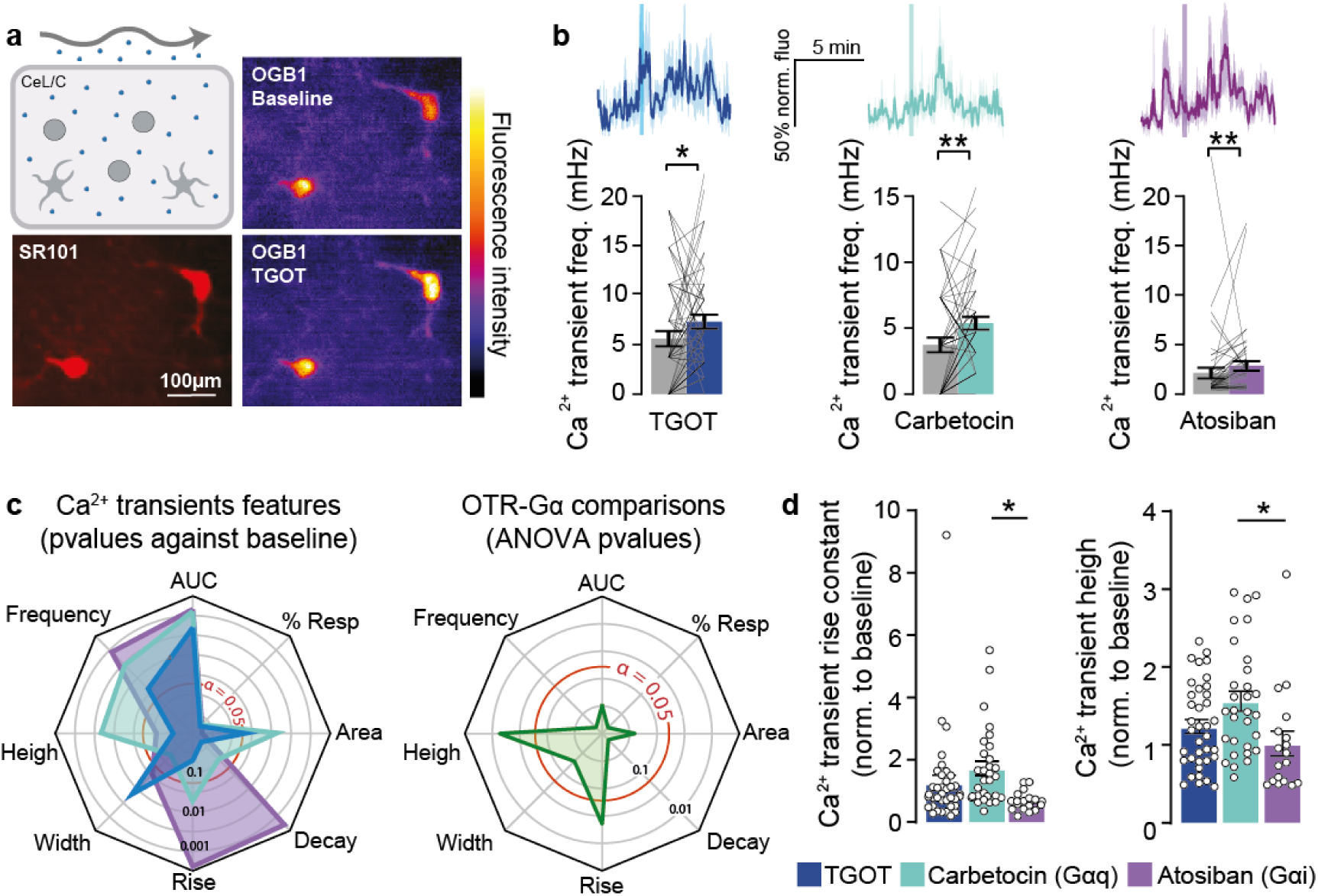
Astrocytic intracellular pathway activated downstream OTR. **A.** Calcium imaging of SR101-identified CeL/C astrocyte somata in brain slices before and after bath application of OTR agonists. Micrograph of slice during patch-clamp experiment. Scale bar=100µm. **B.** Upper panel: mean trace of the normalized to the baseline OGB1 fluorescence recorded in CeL/C astrocytes in response to TGOT (500nM, blue), Carbetocin (500nM, light green), or Atosiban (500nM, blue). The vertical line indicated the start of agonist application. Lower panel: frequency of Ca^2+^ transients in CeL/C astrocytes. n_TGOT_=57 astrocytes over 13 slices, n_atosiban_=49 astrocytes over 9 slices, n_carbetocin_=54 astrocytes over 13 slices. **C.** Comparison of calcium activity triggered by different OTR agonists. Spider plots showing (left) the p-value of the comparison between the basal state and after the application of different agonists for different calcium signaling parameters and (right) the p-value of a one-way ANOVA to compare calcium events triggered by the three different agonists. **D.** Bar plots showing mean rise time and height of calcium transients triggered by indicated OTR agonists normalized to baseline values. Data are expressed as mean across cell ± SEM. Detailed statistics can be found in *Statistic Table 6*. * p<0.05, ** p<0.01, *** p<0.001.

### OTR-Gαi signaling mediates CeL/C astrocytes morphological reorganization and controls neuron sensitivity to extracellular K^+^

Given the involvement of OTR-Gαi in CeL/C astrocytes Ca^2+^ activity, we sought to elucidate the potential role of OTR-Gαi-mediated Ca^2+^ transients in CeL/C astrocytes. To do so, we incubated brain slices with different OTR agonists for 1h and assessed astrocyte GFAP levels by histology (**Fig. 6a**). While the full (TGOT) and Gαi- biased (atosiban) agonists increased CeL/C astrocytes GFAP volume (p_TGOT_=0.007, p_atosiban_<0.001), the Gαq-biased agonist (carbetocin) did not (p_carbetocin_=0.404) (**Fig. 6b**). This suggested an involvement of Gαi, but not Gαq, in the modification of the GFAP cytoskeleton. Similarly, TGOT increased the K^+^-induced depolarization of CeL/C neurons (p=0.003), an effect prevented by the co-incubation of TGOT with the OTR antagonist dOVT (p>0.999) or by genetic deletion of OTR (GFAP OTR cKO, p>0.999) (**Fig. 6c-d** and **Extended data Fig. 6a-d**). These results indicate that astrocytic OTR-Gαi is mediates the OT-induced hypersensitivity of CeL/C neurons to extracellular K^+^. Similarly, atosiban but not carbetocin increased the K^+^-evoked depolarization of CeL/C neurons (p_atosiban_=0.022, p_carbetocin_>0.999; **Fig. 6d** and **Extended data Fig. 6a-d**).

**Figure 6:**
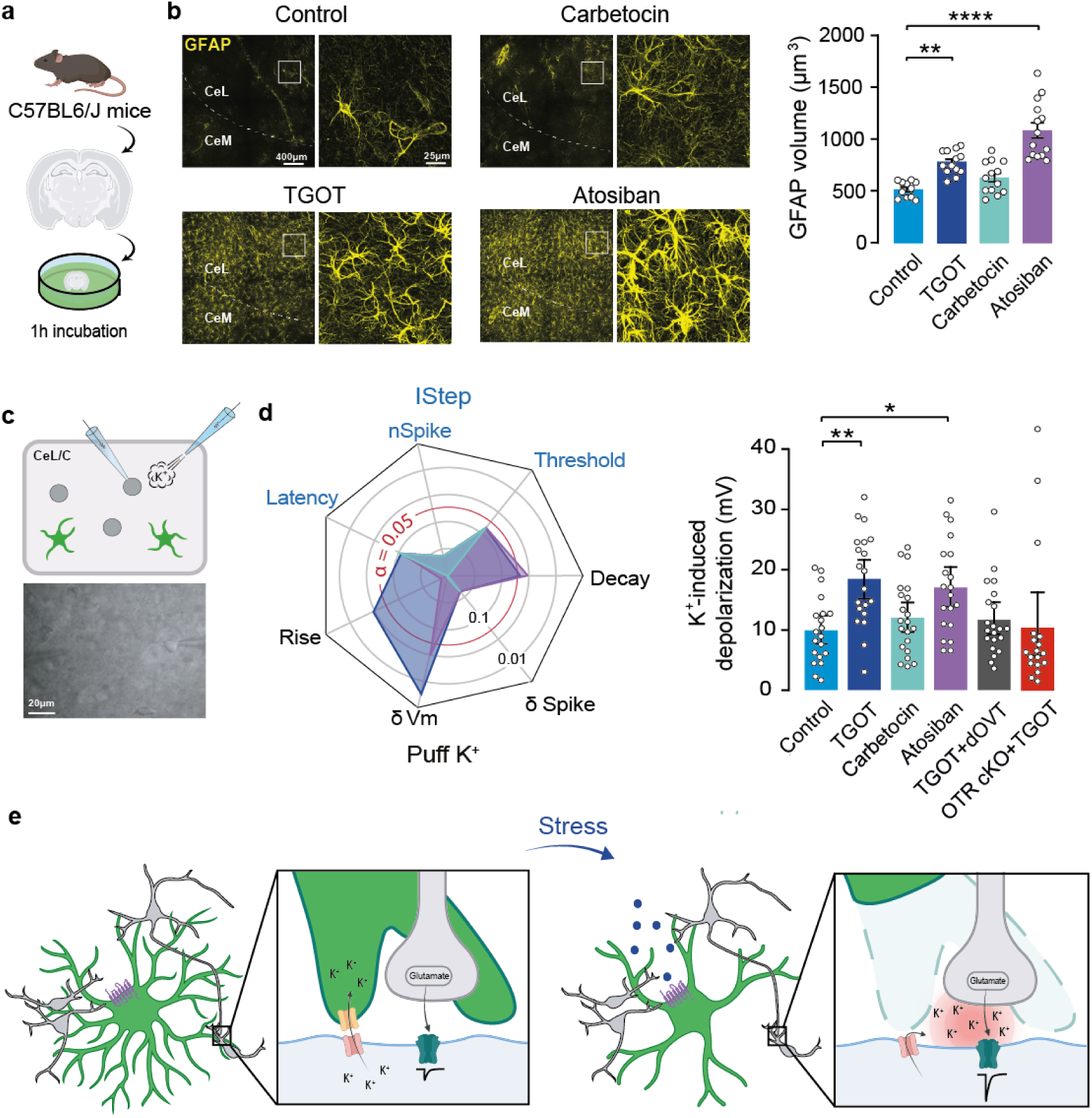
Effect of the different OTR intracellular coupling on the astro-neuronal network. **a.** Analysis of OTR-Gαi activation on CeL/C astro-neuronal network. Brain slices were incubated for 1h in the different OTR agonists before fixation and GFAP staining. **b.** Left: Representative micrograph showing GFAP staining of CeL/C astrocytes in presence of indicated agonists. Scale bars=400 and 25µm. Right: Mean values of the GFAP volume in CeL/C astrocytes under indicated conditions. n_Control_=134, n_TGOT_=208, n_atosiban_=271, n_carbetocin_=225 astrocytes. **c.** Electrophysiological recordings from CeL/C neurons following OTR agonist application. Micrograph of slice during patch-clamp experiment. Scale bar=20µm. **d.** Left: Spider showing the p-values given by the comparison of different electrophysiological features. Current steps were used to measure the spike-firing threshold, the number of spikes at the second active step (nSpike), and the latency to the first spike. K^+^ puffs (30mM in aCSF), were used to measure the difference in spike firing (δSpike), the amplitude of the K^+^-induced depolarization (δVm), and its rise/decay constants. Right: Amplitude of membrane depolarization during the K^+^ puff. n=20-21 neurons. **e**. Model summarizing roles of astrocytes in OTR-mediated stress behavior. Under naive conditions (left) CeL/C astrocytes express few GFAP filaments and buffer potassium at the BLA-to-CeL/C synapse. After acute stress induction (right), astrocytes retract their processes away from synapses and decrease K^+^ buffering leading to an increased strength of BLA-to-CeL/C synapses. Data are expressed as mean across cell ± SEM. Detailed statistics can be found in *Statistic Table 6* and other electrophysiological parameters of CeL/C neurons can be found in *Supplementary* Figure 6. * p<0.05, ** p<0.01, *** p<0.001, **** p<0.0001.

Collectively, these findings suggest that the OTR-Gαi pathway in CeL/C astrocytes recapitulate features of acute stress-induced morphological alterations in astrocytes and adjacent neurons sensitivity to extracellular K^+^ (**Fig. 6e**).

## Discussion

We here provide evidence that acute stress induces a series of OTR- and Ca^2+^ dependent changes in astrocytes of the amygdala, which ultimately impact synaptic transmission in this brain area. Proteomic analysis hinted us towards stress-induced structural remodeling of astrocytes, as verified by three-dimensional reconstruction of astrocytes morphology. Interestingly, OTR-dependent astrocyte plasticity was associated with an increased sensitivity of CeL/C neurons to extracellular K^+^, a decreased in astrocyte synaptic coverage, and an increased strength of BLA-to-CeL/C synapses. Crucially, this morpho-functional plasticity depends upon the recruitment of OTR-Gαi pathway in astrocytes. We propose that this mechanism is key to achieve anticipated behavioral response to a potential danger, contributing to animal survival.

Previous studies have shown that activation of OTR in the CeA decreases freezing behavior induced by Pavlovian fear conditioning in rodents during fear expression and extinction, but not during fear acquisition^8,26,35^. In contrast, in the context of acute stress, GFAP OTR cKO mice do not exhibit freezing behavior, indicating that astrocytic OTR-mediated signaling is necessary for this response. The acute stress paradigm used in our study resemble to a fear acquisition protocol, where a conditional stimulus (sound) precedes an unconditional stimulus (shock). This suggests that astrocytic OTR in the CeA is crucial not only for stress-induced freezing but also for fear acquisition. Therefore, the OTR’s roles in freezing behavior may stem from the varying effects of OT depending on the physiological state of the animal^36^. In summary, OT enables freezing in response to a potential danger during acute stress, while it may trigger a fight-or-flight response when facing a known danger during fear. The physiological relevance of OT, and its action on astrocytes, is then of critical importance: it enables the animal to execute the appropriate behavior in response to a potential threat.

A cardinal feature of the cellular response to acute stress was the retraction of astrocytic processes leading to reduced coverage of CeL/C synapses. This type of structural plasticity has also been observed in the lateral amygdala, where fear conditioning induce the retraction of astrocytic processes from synapses^28^. Interestingly, similar OT-dependent structural changes were also observed in the hypothalamus after prolonged stimulations such as lactation^28,37–39^. These results suggest that altering their morphology is a general mechanism by which astrocytes respond to disruption of brain homeostasis. Of importance, OTR removal from CeA astrocytes prevented stress-induced freezing in male, but not in female mice, suggesting potential sex-specific mechanisms. Sex differences in the OT/OTR system have been reported^40,41^, with female mice having more OTR in the CeA as compared to males^42^. The described inverse correlation between GFAP levels and branching of astrocytes resembles the reactive astrocytes state, where essential physiological functions such as K^+^ buffering are altered^43^. These include the upregulation of the transcription factor STAT3 and of intermediate filaments vimentin and GFAP. Furthermore, amygdala astrocytes show an acute-stress induced downregulation of K_ir_4.1 protein expression, also observed in some pathological contexts. Astrocytes can rapidly modify their morphology upon strong neuronal activity in calcium-dependent manner^44,45^. In accordance with our findings, it was reported that Gαi pathways activate the transcription factor STAT^46^, which is a key mediator of astrocyte reactivity in many pathological contexts^47^. Moreover, two recent studies linked Gαi-GPCR activation and astrocyte morphology^48,49^, indicating this pathway to be a potentially common mechanism for the regulation of astrocyte morphology. Especially, Soto and colleagues showed a decreased astrocyte morphology and homeostatic functions in mice from a model of compulsive and anxiety-related behaviors. Chemogenetic stimulation of the Gαi pathway in astrocytes reverse these phenotypes^49^. Thus, it is tempting to speculate that astrocytic Gαi proteins evoke intracellular Ca^2+^ transients that subsequently modify astrocyte morphology during demanding physiological and behavioral challenges, whose dysregulation might be linked to pathological conditions.

Remarkably, both OTR-Gαi and OTR-Gαq pathway activation led to Ca^2+^ transients in astrocytes, which induced distinct cellular responses. How do astrocytes differentiate between these two signaling pathways to trigger an appropriate reaction? It seems plausible that concomitant activation of Gαq and Gαi signaling pathways may occur upon OTR activation. To support this hypothesis, we observed that albeit Gαi blocking by pertussis toxin prevent atosiban but not TGOT-induced Ca^2+^ signaling, Gαq blocking prevents astrocytes response to both carbetocin and TGOT, indicating that TGOT-induced Ca^2+^ response does not rely on Gαi activation only. This can be explained by the property of Gβγ subunits of the Gαi protein to enhance subthreshold Gαq signaling^50^, suggesting that a concomitant activation of these proteins is needed to generate Ca^2+^ transients. Such signal may encode a specific message, as suggested by the distinct features in the Ca^2+^ events triggered by astrocytic Gαi- or Gαq-OTR pathways. This discovery suggests that these pathways likely exert differential effects on cellular functions such as gliotransmission or morpho-functional changes. In line with this, our results indicate that an OT-induced astrocyte morphological change only occurs upon activation of OTR-Gαi, but not Gαq, pathway. This finding is supported by recent studies, which elegantly show that whilst Gαi-GPCR signaling is linked with changes in astrocyte shape^48^, Gαq-GPCR activation does not regulate GFAP expression^20^. This suggests that Gαi pathway is a potentially common signaling involved in the regulation of astrocyte morphology. A striking feature of our observations is the short amount of time needed to establish cellular and molecular changes following stress exposure. Astrocytes’ shape functions modifications are frequently described in pathological contexts^43^, however, such changes in physiological contexts remains unexplored. In the present study, we used a 10 minutes stress session followed by 5-10 minutes to anesthetize the mice and perform the slices or fixate the tissues. The morphological and homeostatic changes that occurs within 15-20 minutes can seems surprisingly fast but is supported by live imaging of astrocyte processes where a significant retraction can be observed after 10-20 minutes following LTP induction^44^. Finally, proteomic analysis of amygdala samples after acute stress showed profound changes of the protein networks with an unexpected range and magnitude, given the short duration of stress exposure (10min). This data revealed a strong signature for altered astrocyte functionality across various classes of proteins, including channels, transporters, receptors, intermediate filaments and Gαi pathway. This broad coverage was afforded by the deep proteome profiling spanning >8000 proteins, which, to the best of our knowledge, represents the largest amygdala proteome dataset to date. While we focused on astrocyte-specific stress response proteins, these data will provide a rich resource for further mining by the community.

In conclusion, our study suggests that acute stress induces OTR-dependent astrocyte morpho-functional plasticity associated with a decrease in synaptic coverage and an increased strength of BLA-to-CeL/C synapse. Together, this novel mechanism may represents the basis of proper anticipated behavioral response to stressful situations, contributing to mammalian survival.

## Methods

### Animals

All experiments were conducted in accordance with European Union rules and approbation from the French Ministry of Research (01597.05). For *ex vivo* and *in vivo* experiments, male and female C57BL/6j mice were used. For e*x vivo* experiments animals were between 1 and 3 months old. To target GCaMP6f to OT neurons in the PVN, Ai148 mice (Jackson Laboratory, 030328) were crossbred with OXT-Cre (Jackson Laboratory, 024234). Animals were housed under standard conditions with food and water available *ad libitum* and maintained on a 12h light/dark cycle.

### Viral transduction following stereotaxic injections

Animals were deeply anesthetized with 4% isoflurane, and their heads were fixed in a KOPF (model 955) stereotaxic frame. Metacam (Meloxicam, 20mg/kg) was injected subcutaneously to limit inflammation and local analgesics were injected locally at the incision site (Bupivacaïne, 5mg/kg, and Lurocaïne, 2mg/kg). The skull was then exposed, and two holes were drilled according to coordinates adapted from Paxinos and Watson brain Atlas. For injections in the CeL/C, we injected 200nL of the appropriate rAAV at the following coordinates: rostrocaudal = -1.25mm, mediolateral = ±3.1mm, dorsoventral = -5.0mm. To stimulate BLA neurons, we injected 100nL of AAV9-CaMKIIa- hChR2(H134)-EYFP at the following coordinates: rostrocaudal = -1.8mm, mediolateral = ±3.4mm, dorsoventral = -5.0mm. The wound was then sutured and the weight of the animals was followed for five days to verify the post-operatory recuperation. After ∼3 weeks, mice were used for experiments.

### Behavior analysis

***Stress protocol.*** The day before the stress protocol, animals were introduced in the stress box with a smooth white plastic floor and walls for ten minutes for habituation and internal control. Mice were exposed five times to a 20s-long CS tone (2kHz, 60dB) with a variable inter-tone interval (20 to 180 seconds). The next day, stripe patterns were placed on the walls of the box and the smooth white plastic floor was removed, revealing the grid floor of the box. Animals were re-introduced in the stress box for 10min and exposed to five tones with different inter-tone time intervals. The last second of these tones coincided with a 1s-long foot shock of 0.6mA. Freezing behavior, speed and distance traveled was quantified using the ANY-maze software. Freezing was defined as periods longer than 2 seconds where mice did not move and was measured during the 20s sound exposition. For subsequent *ex vivo* experiments, mice were injected with anesthetics directly after the end of the stress protocol and slices were performed in the following 10 to 20 minutes.

***Sucrose preference test.*** Cages were equipped with two feeding bottles, one with sucrose 1% and the other drinking water. At 24h after the beginning of the experiment, we reversed the position of the bottles to avoid a potential place preference. Bottles were weighted before the experiment, 24h and 48h after. We then calculated the sucrose preference according to the following formula: preference = (sucrose intake/total intake) × 100.

***Conditioned place preference.*** In this test, animals develop a preference to a clonidine-paired chamber due to both pain relief in this environment and avoidance for the saline-paired chamber associated with ongoing pain. The apparatus (42553 Conditioned Place Preference for Mice, Ugo Basile, Gemonio, Italy) consists of 2 Plexiglas chambers separated by manually operated doors. Two chambers (size 16 x 25 x 25cm) distinguished by the texture of the floor and by the wall patterns. On the first, second and third days (pre-conditioning), animals were free to explore the apparatus during 30min, and the time spent in each chamber was recorded using ANY-maze™ software. Animals spending more than 75% or less than 25% of the total time in one chamber were excluded from the study. On the 4^th^ day (conditioning) animals were intrathecally injected with either saline solution (10µL, morning) or clonidine solution (10µg in 10µL in the afternoon) and restricted in one chamber for 15min, switching chamber between the morning and the afternoon. On the 5^th^ day the animals were free to explore the 3 chambers and the time spent in each chamber was recorded for 30.

***Von Frey test.*** The mechanical threshold of hind paw withdrawal was evaluated using von Frey hairs (Aniphy Vivo-Tech, Salon-de-Provence, France). Mice were placed in clear Plexiglas® boxes on an elevated mesh screen. After a habituation time of around 10min, the filaments were pressed on the plantar surface of each hind paw in a series of ascending forces (0.6 to 8 grams). Each filament was applied five times per paw, until it just bent, and the threshold was defined as three or more withdrawals observed out of the 5 trials. The mean of sensitivity of the two hind paws was calculated.

***Hargreaves test.*** The latency for hind paw withdrawal in response to thermal stimulation was determined using the Hargreaves method. Mice were placed in clear Plexiglas® boxes and testing began after exploration and grooming behaviors ended (15min). The infrared beam of the radiant heat source (37570 Plantar Test, Ugo Basile, Gemonio, Italy) was applied to the plantar surface of each hind paw. The experimental cut-off to prevent damage to the skin was set at 15s. Three measures of the paw withdrawal latency were obtained per animal and averaged for each hind paw.

***Dry ice test.*** Hind paw withdrawal latency in response to cold thermal stimulation was determined using the dry ice method. Mice were placed in clear Plexiglas® boxes. After a 15min habituation period, a cut 2mL syringe filled with dry ice powder was placed against the glass surface under the animal’s paws. This source of cold under the animal’s plantar surface caused the animal’s paw to retract. Latency times were recorded manually using a stopwatch. Three measurements of paw withdrawal latency (spaced at least 15min apart) are then taken and averaged for each hind paw.

***Social interaction test.*** Social interactions were assessed in the cage of the same size of the home cage, with a bit of dust at the bottom. For this test, interactions between a test subject and a stimulus mouse were systematically scored and videotaped for 5min. Light intensity was 150lux. All cages were bring in the test room for acclimatization of 30min. Then, the stimulus and test mice were introduced in the same time in a clean cage for 5min. The stimulus mice were from the same background C57BL6, same sex, same age and used maximum twice a day. Their weight was on average 7% less compared to test mice. Order of testing was counterbalanced according to genotype and stimulus mice. The times the test subject spent performing social behaviors (i.e. facial sniffing (oral-to oral contact), anogenital sniffing (oral-to-base sniffing), direct touching with a paw and pursuit of the stimulus mouse) were determined in real time with an ethological keyboard (ANY-maze software). After testing, each mouse was returned to its home cage and a new clean cage was used for the next mice. The percent of time spent engaged in social behavior compared to total time (5min) was calculated for the measure of social interactions:

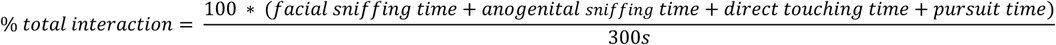

### Corticosterone measurements in blood

***Sampling.*** Blood sample (40μl) were collected from tail tips following a scissor incision using a calibrated sodium-heparinized capillary (40µl, Hirschmann™ Disposable Capillary Tubes, Fisher Scientific, Illkirch, France). The blood was then supplemented with 10µl of heparin (843 µM; Sigma-Aldrich, St. Quentin Fallavier, France). Subsequently, 10µl of D4-CORT (5µM, Sigma-Aldrich) in 99.1% H_2_O/0.1% formic acid (v/v; Fisher Scientific) was added to the mixture. Proteins were precipitated acetonitrile (ACN 100%, Fisher Scientific) and samples were centrifuged at 20,000g (4°C, 30min). The supernatant was collected and dried under vacuum (SpeedVac, Thermo Fisher). The dried samples were then resuspended in 20μl of 20% ACN/0.1% formic acid (v/v). The supernatants was collected and stored at -80°C.

***Dosage.*** Corticosterone measurements were conducted using a Dionex Ultimate 3000 HPLC system (Thermo Electron, Villebon-sur-Yvette, France) coupled to a triple quadrupole Endura mass spectrometer (Thermo Electron). The samples were loaded onto a heated Hypersil aQ column (100 x 1mm, 1.9μm, with a flow rate of 90μl/min; Thermo Electron) at 40°C. Qualification and quantification were performed using multiple reaction monitoring (MRM) based on the isotopic dilution method. Compound identification was achieved by comparing the precursor ions, selective fragment ions, and retention times with those of the isotopically labeled standards (IS). The selection of monitored transitions, along with the optimization of collision energy and RF lens parameters, was automated (*Supplementary Data Table “HPLC and MSMS details”*). System control was driven by Xcalibur v4.0 software (Thermo Electron).

### Microdialysis

Male CD1 mice (25-30g; 7-8 weeks old; Charles River, Germany) were stereotaxically implanted with a U-shaped microdialysis probe (molecular cut-off: 18kDa; length of membrane 1.5mm) under isoflurane anaesthesia (Isoflurane Baxter, Baxter Deutschland GmbH, Germany) targeting the right CeA (AP: -1.0; lateral +2.4; DV: -5.0 mm from bregma; Paxinos and Watson, 1998). After a 48h post-surgery recovery, the microdialysis probe was first perfused without sampling for 90min (3.3µl/min, Ringer’s solution, adjusted to pH 7.4), before 3 consecutive 30min microdialysates from the CeA were collected (1) under basal conditions, (2) during exposure to 5 random foot-shocks (0.6mA, 1s) over a 10min period in a conditioning chamber (45 x 22 x 40cm, transparent Perspex box with a steel grid floor), and (3) again in the home cage. Samples were kept at -20°C until radioimmunological analysis of OXT (detection limit: 0.5pg/sample, intra-assay variability of <8% cross-reactivity with other neuropeptides including arginine vasopressin <0.7%). Brains were frozen and the perfusion sites were histologically verified on 40µm cryo-cut stained brain slices.

### In vivo imaging

***In vivo micro-endoscope calcium imaging.*** To visualize calcium level in PVN OT neurons, Ai148:OXT-Cre mice were stereotactically implanted with a chronic GRIN lens (0.5mm in diameter; 7.4mm in length, Inscopix) ± 200µm above the PVN at the following coordinates: rostrocaudal = - 0.85mm, mediolateral = +1.1mm, dorsoventral = -4.9mm with a 10-degree angle (right hemisphere). The lens was securely fixed to the skull with a three-component dental cement (C&B-Metabond, Parkell Inc.). After a recovery period of at least two weeks, animals were implanted with a baseplate to attach the micro-endoscope (nVista 3, Inscopix). A tethered dummy micro-endoscope was connected to the baseplate for habituation to the recording procedures. Following an additional week for recovery and habituation, the real miniature microscope was attached to the baseplate one day before the experiment. Imaging data were collected at 10Hz, with excitation light power 0.8-1.2mW/mm² (40-60% of maximal power, nVista 3, Inscopix). Stacks of calcium images were first preprocessed to correct time-invariant pixels or reconstruct frames lost during the acquisition, and were then adjusted for motion artifacts in the x-y plans using Inscopix Data Processing software 1.9.1 (Inscopix). Subsequently, image stacks were exported to ImageJ, where the recording background was subtracted. Regions of interest (ROI) corresponding to individual neurons were manually defined over z-stacked images, and the raw average fluorescence from each ROI was quantified and exported using Fiji software. Raw traces are further processed and normalize using custom-made scripts (Matlab, MathWorks). Due to the potential bleaching of the GCaMP6f fluorescence within the session, an exponential curve was fitted and subtracted from the raw signal of each cell during each session to ensuring consistency in the fluorescence measurements. Subsequently, F0 was determined using Gaussian mixture modeling, where the observed intensity values from each cell in each session were approximated as a mixture of Gaussian distributions via an expectation-maximization procedure. The mean of the distribution containing the lowest values was identified as the F0 for each neuron and session. ΔF/F0 are calculated base on bleaching-corrected calcium traces and the estimated F0, and then are normalized using envelope normalization.

***Fiber photometry.*** 3-4 weeks after viral injection, fiber-optic cannula (fiber core, 400μm; 0.37 NA; 5 or 5.5mm length; Doric Lenses, Quebec City, Canada) was implanted approximately 0.1mm above the injection site and fixed to the skull with dental cement and Super-Bond C&B. After 5-7 days recovery, mice were handled by the experimenter for 3-5 days and acclimated to the fiber-optic patch cord connection for 3-5 additional days. Fluorescence intensity was measured with a dual-color fiber photometry setup equipped with two LED light sources, 465 and 560nm (Doric Lenses). The light intensity at the end of fiber-optic patch cord was adjusted to between 50 and 80µW. Data were acquired by lock-in mode in Doric Neuroscience Studio. Acquired data were processed by custom-made python codes. After data denoising with a Savitzky-Golay filter, photobleaching was corrected for by a computed control channel generated by fitting with an exponential curve with the corresponding signal. Normalized ΔF (ΔF/F%) was calculated by the equation: 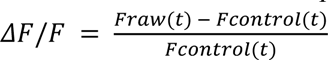 where Fraw is the Ca^2+^-dependent fluorescence changes and F_control_ is the value in control channel computed for minimizing the photobleaching effect.

### Proteomic analysis

***Tissue sample preparation.*** Ten to twenty minutes following the end of the stress protocol, mouse brains were sliced as described for *ex vivo* electrophysiology. Using a 1mm-diameter punch, the CeA was dissected bilaterally on two 400µm slices per hemisphere (4 punches per mouse), snap-frozen and stored at -80°C. Isolated amygdala tissues from naïve and stressed mice (n=5 per group) were disrupted using a lysis solution containing 1% Sodium-dodecylsulfate, 50mM Tris (pH 8.0), and a protease inhibitor cocktail (Merck) without EDTA. The tissues were subjected to sonication in AFA-tube TPX strips (Covaris) using a Covaris LE220R-Plus instrument (300s, average power of 162.5, peak power of 325, 50% duty factor, 200 cycles per burst). Tolerance of the dithering settings were set at ± 5mm in the x and z axis, and 4.5mm in the y axis, while operating at a velocity of 20mm/s. Protein concentrations of the sonicated lysates were measured using a BCA assay (Pierce). For mass spectrometer analysis, protein digestion and peptide generation were performed using 1μg of protein per condition. Due to the limited amount of initial material, we utilized the automated version of the SP3 protocol for protein digestion and peptide purification. The Bravo system is designed to handle 96 samples at once, performing all necessary steps such as protein reduction and alkylation, magnetic bead aliquoting, protein clean-up using SP3, protein digestion, and peptide recovery, as described in detail before.

***LC-MS/MS.*** The quantification of tryptic peptides from the whole proteome was performed using an EASY-nLC 1200 system (Thermo Fischer Scientific) connected to a Fusion mass spectrometer (Thermo Fischer Scientific). Peptides were separated by reverse-phase liquid chromatography employing 0.1% formic acid (solvent A) and 80% acetonitrile (solvent B) as the mobile phases. The separation of peptides was performed using an Acclaim PepMap trap column (Thermo Fisher Scientific, C18, 20mm × 100μm, 5μm C18 particles, 100Å pore size) and a nanoEase M/Z peptide BEH C18 analytical column (Waters, 250mm × 75μm 1/PK, 130Å, 1.7μm). The gradient started with a 3% concentration of solvent B for the first 4min, thereafter increasing to 8% and further rising to 10% after 6min. The concentration of solvent B increased to 32% after 68min and to 50% after 86min. Between the 87th and 94th minute of the gradient, the concentration of solvent B reached 100%. The system was re-equilibrated after 95min using 3% of solvent B for 10min. The eluted peptides were subjected to ionization and introduced into the mass spectrometer using the Nanospray flex ion source (Thermo Fischer Scientific) and a Sharp Singularity nESI emitter (ID = 20μm, OD = 365μm, L = 7cm, α = 7.5°) (FOSSILIONTECH), which was connected to a SIMPLE LINK UNO-32 (FOSSILIONTECH). A constant spray voltage of 2.5kV was applied to the emitter, and the capillary temperature of the ion transfer tube was set to 275 °C. The Fusion mass spectrometer was used in data-independent (DIA) mode with a scan range of 400-1000m/z, an Orbitrap resolution of 60000 FWHM, an AGC target of 3e6, and a maximum injection duration of 20ms. The acquisition of data-independent MS/MS spectra was performed at a resolution of 30000 FWHM. The maximum injection duration was set to 50ms and the AGC target was set to 1e6. The acquisition strategy included employing 26 isolation windows, each having a width of 23.3m/z and overlapping by 1Th. The collision energy used for fragmenting the precursor ions was set to 27, and a fixed initial mass of 200m/z was used to get the MSMS spectra.

***Data analysis.*** Raw data obtained from DIA measurements were analyzed using DIA-NN version 1.8.0. A spectral library was developed based on the predictions made by the fasta file. In addition, a fasta file containing commonly seen protein contaminant was included for the purpose of predicting the spectral library. The spectral library prediction used the default values. The default parameters of DIA-NN were used for the processing of the raw files. The precursor matrix output tables were filtered to include only values with a false discovery rate <0.01. Additionally, values were filtered based on a channel q-610 value <0.01 and a translated q-value <0.01. MBR feature was activated to enhance data integrity. Output tables generated by DIA-NN were then analyzed with R-Studio, using the dplyr package for data wrangling, and ggplot2 for visualisation. Differential expression and statistical analyses were carried out using the limma package. Gene Set Enrichment Analysis (GSEA) was conducted with the fgsea package, heat maps were created using the pheatmap package and Perseus software.

### Immunofluorecence staining and image analysis

***Tissue fixation.*** Mice were anesthetized using ketamine (Ketamin 1000, 400mg/kg) and xylazine (Paxman, 80mg/kg) administered intraperitoneally. Animals were then perfused transcardially with PBS, followed by 4% PFA. Brains were dissected and post-fixed overnight in 4% PFA at 4°C.

***Astrocyte morphology assessment.*** 50μm vibratome sections were cut, collected, and stored in PBS 1X overnight at 4°C. Slices were then incubated in a PBS1X-Gelatin 0.02%-Triton 0.25% solution for permeabilization and saturation of non-specific sites during 1h at RT. Incubation of the first antibodies took place overnight at 4°C and slices were then washed and incubated with secondary antibodies overnight at 4°C. Finally, slices were mounted in an Abberior antifade medium and left for one day at 4°C in the dark for polymerization. All antibodies used and their concentrations are given in the reagent table (*Supplementary Data Table “List of reagents”*). Z-stack images of the CeL were captured (6-8 images per animal) with a depth of 50μm and 1μm intervals, employing a 40x magnification on the Nikon AX confocal system (Nikon Imaging Center, Heidelberg). All images used for analysis were taken with the same confocal settings (pinhole, laser intensity, digital gain, and digital offset). Imaris software (version 10.0, Oxford Instruments) was used for reconstruction and morphological analysis of astrocytes. Raw confocal files were imported into Imaris and subjected to surface reconstruction, involving background subtraction and setting surface detail to 0.5µm with the largest diameter at 0.5µm. Immunofluorescent signals from GFAP staining or virally-expressed GFP were utilized as templates for surface and filament reconstructions. Astrocytes falling below a volume of <250µm³ or exceeding >5500µm³ were excluded from data analysis. A separate masked channel was generated for filament analysis, employing automated threshold detection with maximum seed point placements. Manual intervention ensured each astrocyte possessed precisely one starting point at the soma center. Morphological analysis involved calculating average values for each animal individually (surface area, volume, etc.), with data points representing mean values for each morphological parameter. Sholl analysis was conducted using Imaris in the filament reconstruction mode, and data sets were to Excel files for further analysis. Each data point on the respective Sholl plots corresponds to the mean Sholl intersections, accounting for the average of the respective cohort with error bars indicating ±SEM, calculated across all astrocytes per animal. For synaptic engulfment analysis, object-to-object statistics were activated, enabling quantification of engulfed Homer1 or vGlut1/2 based on the surface-to-volume ratio. Synaptic compartments were reconstructed using specific settings including background subtraction, surface detail at 0.2µm, and the largest diameter at 0.2µm. Debris measuring <0.5µm³ was disregarded as nonspecific and filtered out in a subsequent step.

### Ex vivo recordings

***Slice preparations.*** Mice were anesthetized using ketamine (Ketamine 1000, 400mg/kg) and xylazine (Paxman, 80mg/kg) administered intraperitoneally. Intracardiac perfusion was then performed using an ice-cold NMDG-based aCSF containing (in mM): NMDG (93), KCl (2.5), NaH2PO4 (1.25), NaHCO3 (30), MgSO4 (10), CaCl2 (0.5), HEPES (20), D-glucose (25), L-ascorbic acid (5), thiourea (2), sodium pyruvate (3), N-acetyl-L-cysteine (10) and kynurenic acid (2). This solution was bubbled in 95% O_2_/5% CO_2_ gas all the time of the experiment and its pH was adjusted to 7.3-7.4 using HCl. After decapitation, the brain was swiftly removed in the same ice-cold dissection aCSFs and hemisectionned. 350μm-thick horizontal slices containing the CeA were obtained using a Leica VT1000S vibratome. Slices were then stored in 37°C NMDG aCSF for 10min before placing them in the holding chamber at room temperature containing normal aCSFs for 1 h minimum before any experiments were conducted. Normal aCSF used for all *ex vivo* experiments was composed of (in mM): NaCl (124), KCl (2.5), NaH_2_PO_4_ (1.25), NaHCO_3_ (26), MgSO_4_ (2), CaCl_2_ (2), D-glucose (15), adjusted for pH values of 7.3-7.4 with HCl or NaOH and continuously bubbled in 95% O_2_/5% CO_2_ gas. Osmolality was set to 300±10 mOsm/L. For pharmacologic experiments, slices were incubated in aCSF supplemented with 500nM of agonist and/or antagonist for 1h. Slices were then transferred to an immersion-recording chamber superfused at 2ml/min with aCSF.

***Electrophysiology.*** Whole-cell patch-clamp recordings of CeL/C neurons were done using infrared oblique light visualization. For astrocyte recordings, slices were incubated in 1µM SR101 at 37°C for 20min and washed for 1h. Subsequent patch-clamp recordings from astrocytes were guided by SR101 fluorescence. Borosilicate glass electrodes (OD 1.5mm, ID 0.86mm; Sutter Instrument) were pulled using a horizontal flaming/brown micropipette puller (P97; Sutter Instrument) to obtain 3.5-7 MΩ recording pipettes. These pipettes were filled with an intracellular solution containing (in mM): K-Glu (125), HEPES (20), NaCl (10), and ATPNa_2_ (3). The pH was adjusted to 7.3-7.4 with KOH, and osmolality was adjusted at 300±10 mOsm/L with water. Recordings were acquired with an Axon MultiClamp 700B Amplifier coupled to a Digidata 1440A Digitizer (Molecular Devices) at 20kHz using either the pClamp 10 software suite for puff experiments or using WinWCP 4.2.2 freeware (John Dempster, SIPBS, University of Strathclyde, UK) for photostimulation experiments. Series resistance was monitored and manually compensated (∼50% typically). At the beginning of each recording, current pulses ranging from -10pA to 100pA with 10pA steps was applied for 300ms per sweep to neurons or voltage pulses ranging from -100mV to 100mV with 20mV steps lasting 100ms for astrocytes. For neurons, the membrane potential drift was estimated with the Savitsky-Golay polynomial local filter from the scipy library and subtracted from the membrane potential. Spikes were then detected with the find_peaks function of the Scipy library for each sweep as peaks with a height and a prominence higher than 10mV. The latency of spikes was quantified as the time between the beginnings of the first step that trigger a spike and the first spike. The number of spike was measured in the second step that triggers action potentials. This protocol was repeated three times and the median results were used for further analysis. For astrocytes, voltage-current curve were constructed by measuring the current needed to reach the command voltage during the 25 last milliseconds of the voltage step. For K^+^ puff application, the injected current was adapted to hold the neuronal membrane at -60mV. Basal activity of neurons was recorded for 20s and then a 20s long puff a 30mM-containing KCl aCSF was done at ∼100µm of the recorded neuron. The membrane potential drift was estimated after a stringent Gaussian filter from the pyABF library, thereby avoiding overestimation of this value due to action potentials. This drift was then subtracted from original traces and spikes were detected with the find_peaks function of the Scipy library as peaks with a height and a prominence higher than 20mV. Spikes and membrane potential were measured in the 20s baseline, the 20s K^+^ puff application and the 20s washout. The sum of spikes and the median of membrane potential were computed over these three periods. For depolarization kinetics, linear functions were fitted to the five first seconds of the rising/decay phase of the K^+^-induced depolarization on the Gaussian-filtered trace. This protocol was repeated three times and the median values of measures were used for statistical analysis. For optostimulation of BLA neurons, the neuronal membrane potential was held in voltage clamp at -60mV. Basal activity was recorded for 20s of basal activity before stimulating glutamatergic BLA neurons with 5ms blue light pulses at 20Hz for 10s. For each recording, CPSE were detected with the find_peaks function of the Scipy library as peaks higher than three time the standard deviation estimated during the 20s baseline. Frequency and amplitude of these CPSE were measured in five seconds-long quantification windows taken just before the photostimulation, during the five first seconds of the photostimulation, and at the end of the recording. After each light pulse, we estimated a failure as the absence of evoked CPSE, or we estimated the rise and decay of the evoked CPSE. Rise and decay constants were measured on exponential and linear fits as the time needed to go from 10 to 90% or from 90 to 10% of the CPSE amplitude, respectively. This recording was repeated three times and the medial value of the three recordings was used for statistical analysis. For K^+^ currents recordings, astrocytes were patch-clamped at -80mV and basal membrane current was recorded during 20s before stimulating glutamatergic BLA neurons with 5ms blue light pulses at 20Hz for 10s. Subsequently, recordings were performed in the presence of Ba^2+^ to block K currents and BLA neurons were stimulated again after 15min. Evoked currents were estimated as the minimum value during the photostimulation subtracted from the minimum value during baseline. The K^+^ current was calculated by subtraction of currents evoked in the presence of Ba^2+^

***Ex vivo calcium imaging*.** After SR101 incubation, the synthetic calcium indicator OGB1-AM was bulk loaded as previously described^51^, reaching final concentrations of 0.0025% (20μM) for OGB1-AM, 0.002% for Cremophor EL, 0.01% for Pluronic F-127 and 0.5% for DMSO in aCSF and incubated for 1h at 37°C Slices were then washed in aCSF for at least 1h before imaging. Only astrocytes co-labeled for SR101 and OGB1 were used. The spinning disk confocal microscope for calcium imaging was composed of a Zeiss Axio examiner microscope with a ×40 water immersion objective (numerical aperture of 1.0), mounted with an X-Light Confocal Unit–CRESTOPT spinning disk. Images were acquired at 2Hz with an optiMOS sCMOS camera (Qimaging). Cells within a confocal plane were illuminated for 20ms at 575nm for SR101 and 80ms at 475nm for OGB1 using a Spectra 7 LUMENCOR. The different hardware elements were synchronized through the MetaFluor 7.8.8.0 software (Molecular Devices). Because astrocytes are mechanosensitive, OTR agonists (500nM) were bath applied for 20s and not puff-applied to avoid mechanical stimulation. All calcium imaging experiments were conducted at room temperature and cells with an unstable baseline were discarded.

All analyses were conducted as described ^51^. Astrocytic calcium levels were measured in manually outlined regions of interest (ROI) comprising the cell body using ImageJ software. Subsequent offline data analysis was performed using a custom-written Python-based script. To take into account slice micro-movements, the SR101 fluorescence values were subtracted from the OGB1 ones. Then, a linear regression was applied to each trace to correct for photobleaching. Calcium transients were detected using the find_peaks function of the SciPy library as fluorescence variation exceeding 8 time the standard deviation and a prominence exceeding it 5 times (for more details, see ^51^). The number of peaks and the area under the curve was quantified before and after the drug application. All data were normalized according to the duration of the recording, and astrocytes were labeled as ‘responsive’ when their AUC or their calcium transient frequency at least doubled after drug application. Because the time after stimulation is longer than the baseline (10min vs. 5min), the probability of observing a spontaneous calcium peak is stronger after stimulation. To avoid this bias, astrocytes with only one calcium peak during the whole recording were not considered responsive. Each calcium transients were then isolated, their area were estimated with the trapezoid method and their duration were measured as their half-maximum full width (HMFW). The rise and decay phases of Ca^2+^ transients were fitted with linear functions and the coefficient of these slopes was used as rise and decay constants. To compare effects of different agonists (Figure S6), evoked Ca^2+^ activity was normalized to the baseline values as ratio (baseline/drug effect).

### BRET experiments in astrocytes

Purifed astrocytes were isolated from P2 C57BL6 mice whole brains by magnetic-activated cell sorting (MACS) (Miltenyi Biotec, Bergisch Gladbach, Germany) with anti-GLAST (ACSA 1; (Jungblut et al., 2012)) MicroBeads according to the manufacturer’s instructions. Cells were plated on poly-L-lysine-coated (Sigma Aldrich, St. Louis, MO, USA) T75 flasks in complete medium, composed by minimal essential medium (MEM, Invitrogen, Life Technologies, Carlsbad, CA, USA) supplemented with 20% fetal bovine serum (FBS) (Gibco, Life Technologies, Carlsbad, CA, USA) and glucose (5.5g/L, Sigma Aldrich, St. Louis, MO, USA). Cells were grown in a humidified incubator at 37°C and 5% CO_2_. Two days before the experiment, astrocytes were harvested with trypsin 0,25% (Gibco, Life Technologies, Carlsbad, CA, USA) and plated in 12-well plates in complete medium at a density to ensure ∼70% confluent cultures at 24h after seeding. Typically, 80,000 cells/cm² surface area of the culture dish were seeded in cell-specific medium. Astrocytes were transfected using Lipofectamine 3000 (Invitrogen, Life Technologies, Carlsbad, CA, USA) according to the manufacturer’s protocol with a DNA to Lipofectamine ratio of 1:3 w/v. A transfection enhancer, the 3000 enhancer reagent (1:2, DNA : Reagent, w/v), was used along with the Lipofectamine 3000 transfection reagent. 1.25µg of plasmids DNA were transferred to each well of the 12-well plates. The cDNA amounts used per well were as following: Gα_x_ Rluc8 (300ng), GFP^10^-Gγ_2_ (250ng), Gβ1 (300ng) and OTR (400ng).

BRET experiments were performed 72h after transfection. Astrocytes were detached with trypsin 0.25% and resuspended in Krebs-Ringer’s-HEPES (KRH) buffer containing (in mM) NaCl (125), KCl (5), MgSO_4_ (1.2), KH_2_PO_4_ (1.2), CaCl_2_ (2), glucose (6) and HEPES-NaOH (25), pH 7.4.

For specific validation experiments, BRET biosensors were expressed in HEK293 cells. To this end, cells were co-transfected using plasmids encoding Gαq-97-Rluc8 or Gαi2-91-Rluc8 and GFP^10^-Gγ_2_, Gβ1, and the OTR. Two days after transfection, cells were washed twice, detached, and resuspended with PBS with 0.5mM MgCl_2_ at room temperature. Cells were distributed on 96-well microplates (100µg of proteins/well) (Optiplate, PerkinElmer Life Sciences) and incubated at room temperature for 5min in the presence or absence of YM (60nM) and two minutes with PBS or OT (10µM).

BRET measurements were performed as previously described using the Infinite F500 multimode microplate reader (Tecan) ^33^. The signals emitted by the Rluc8 donor that allows the sequential integration of light signals was detected with two filter settings (Rluc8 filter, 370-450nm; GFP^10^ filter, 510-540nm). The data were recorded by Infinite F500 after the addition of 5μM Deep Blue C coelenterazine and the BRET signal was calculated as the ratio between GFP^10^ emission and the light emitted by Rluc8 after 2min stimulations with maximal dose of OT (1µM) or vehicle (PBS). All measurements were performed at 37°C. The BRET signal was calculated as the ratio between GFP^10^ emission and the light emitted by Rluc8. The changes in BRET induced by OT were expressed on graphs as “OT-induced changes in BRET ratio” using the following formula:

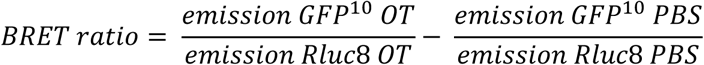

### Tissue preparation, protein extraction and western blotting

Ten to twenty minutes following the end of the acute stress protocol, brains from stressed or naïve (home cage controls) mice were sliced as described for *ex vivo* electrophysiology. Using a 1mm-diameter punch, the CeA was dissected bilaterally on two 400µm slices per hemisphere (4 punches per mouse, tissue weight < 4mg), snap-frozen and stored at -80°C until processing. To enrich membrane protein fraction protein extraction was performed using the Mem-PERTM Plus Membrane Protein Extraction Kit (Thermoscientific) according to the manufacturer’s instructions. Briefly, after rinsing, CeA punches were transferred into grinders (Wheaton; Dounce tissue grinder 1mL, #357538) with 500µL of permeabilization buffer containing protease (1:100, Protease inhibitor cocktail, Sigma-Aldrich, #P8340) and phosphatase (1:100, Phosphatase inhibitor cocktail 2, Sigma-Aldrich, #P5726) inhibitors and grinded until even suspension. Homogenates were transferred in 2mL tubes, incubated for 10min at 4°C under constant rotation followed by centrifugation at 16,000g for 15min at 4°C. After removal of the supernatant containing cytosolic proteins, the pellet was resuspended in 500µL of solubilization buffer with protease and phosphatase inhibitors. Samples were incubated for 30min at 4°C under constant rotation, centrifuged at 16,000g for 15min at 4°C and the supernatant containing membrane proteins was collected and stored at -80°C. Because of very low amounts of input tissue, we used a protein concentration method. Membrane protein samples were incubated with 77% trichloroacetic acid on ice for 30min and centrifuged at 12,000rpm for 15min at 4°C. After supernatant removal, pellets were rinsed with ice cold acetone and centrifuged at 12,000rpm for 5min at 4°C. After supernatant removal, pellets were resuspended in loading buffer (NuPAGE® LDS sample buffer and sample reducing agent, Invitrogen) and denaturated 10min at 70°C. Samples were loaded on 4-20% Criterion™ TGX Stain- Free Precast Gels (Bio-rad) and migration was performed 1h at 170V in Tris-glycine buffer (Bio-rad) followed by Stain-Free gel activation using a ChemiDoc MP Imaging System (Bio-rad). Proteins were transferred on a nitrocellulose membrane with the Trans-Blot Turbo™ Transfer System (Bio-Rad). Total protein load per well was evaluated after imaging the membrane under UV light. After 3x10min rinses in 9mM NaCl Tris-HCl Buffer + 0.1% Tween 20 (TBST), membranes were blocked in TBST + 5% milk for 1h at room temperature (RT) and incubated overnight at 4°C sequentially with the following primary antibodies: anti-Kir4.1 (1:1000, Rabbit ; Alomone, #APC-035), anti-actin or anti-Tom20 in TBST + 5% milk. After 3 x 10min washed in TBST, membranes were incubated for 1h at RT with Horse Radish Peroxidase (HRP)-conjugated secondary antibodies (1:5000, Vector laboratories) diluted in TBST + 5% milk. Membranes were incubated with Clarity Western ECL substrate (Bio-rad) and the signal was detected with ChemiDoc MP Imaging System. Band intensity was quantified with Image Lab Version 5.2.1 and normalized to total loaded protein load. In-between signal detections, peroxidase activity was inactivated using a 20min incubation of membranes in 1% Na-Azide in TBST at RT followed by 3 x 10min wash in TBST. When necessary, antibodies were stripped using a 20min incubation in stripping solution (ReBlot Plus Mild Antibody Stripping Solution, Millipore).

### CeL/C ultrastructural analysis

Mice were anesthetized with intraperitoneal injection of 100mg/kg ketamine chlorhydrate and 5mg/kg xylazine and transcardially perfused with glutaraldehyde (2.5% in 0.1M PBS at pH 7.4). 1mm amygdala punches were immersed in the same fixative overnight, rinsed several times in phosphate buffer, and postfixed in 1% osmium for 1hour. Finally, samples were dehydrated in graded ethanol series and embedded in Embed 812 (EMS). Ultrathin sections were cut with an ultramicrotome (Leica), stained with uranyl acetate (1% (w/v) in 50% ethanol), and examined by transmission electron microscopy (TEM; Hitachi H7500 equipped wuth an AMT Hamamatsu digital camera).

### Statistical analysis

All statictical tests were performed using GraphPad Prism version 8.0.0 (GraphPad Software). Parametrical tests were performed if data showed normal distribution and equality of group variances. Otherwise, non-parametric tests were used. All values, group compositions, and statistical tests for each experiment are detailed in *Supplementary Tables 1-4*.

## Data Availability

The raw data generated in this study are available at public database under accession code (*to be completed upon publication*). All raw proteomic data associated with the manuscript have been deposited to the Pride repository accessible for reviewers via https://www.ebi.ac.uk/pride/login, project number PXD053856, (*Reviewer token 8DIhH2GrahpG*). Statistical data are provided in the *Statistic Tables 1-4*. In addition, all data that support the findings of this study are available from the corresponding authors upon request. Source data are provided with this paper. All the MatLab code for *in vivo* miniscope, Python code for *in vivo* fiber photometry, and for *ex vivo* experiments can be found at in following repository: https://github.com/Team-Charlet/Baudon_et_al.

## Code Availability

All the MatLab code for *in vivo* miniscope, Python code for *in vivo* fiber photometry, and for *ex vivo* experiments can be found at in following repository: https://github.com/Team-Charlet/Baudon_et_al.

## Acknowledgements

This work was supported by the Centre National de la Recherche Scientifique contract UPR3212, the Université de Strasbourg contract UPR3212; the NeuroStra Interdisciplinary Thematic Institute of the ITI 2021-2028 programme; the Agence Nationale de la Recherche (ANR, French Research Foundation) grants n° 23-CE37-0015-01, 19-CE16-0011-0, 19-CE37-0019 and 20-CE18-0031, the FRM Equipe grant EQU202403018071, the Region Grand Est 19_GE9_043 grant (to AC); the Graduate School of Pain EURIDOL, ANR-17-EURE-0022 (to AC, ECC and VGre); the CNRS International Research Project grant ICOT2023 (to AC and VGri); the FRM Fellowship FDT202204015114 (to AB); the Graduate School GRK2174 of the German Research Foundation (to BDB), the EMBO Short Term Fellowship and the Procope Mobility Fellowship (to EV); the graduate school Signaling and Integrated Networks in Biology (to TL); a Fondation Vaincre Alzheimer pilot grant (to LBH); the Deutsche Krebshilfe project 70114190 (to JK); the DFG grants 465/34-1 and 465/36-1 (to IDN), AL 2466/2-1 (to FA) and SFB1158-3 (to VG). We thank Noémie Willem and the ComptOpt plateform (Comportement et Optogénétique, CNRS and CoRTecS Université de Strasbourg) for nociception experiments, the SMPMS plateform for corticostesrone quantification (CNRS and CoRTecS Université de Strasbourg), Carole Escartin for feedback on Western blot experiments, Yuval Podpecan and Moritz Wimmer for assistance with confocal microscopy and image analysis.

## Authors contribution

Project conception, AC; Methodology, AA, AB, AC, AL, FA, JK, K-YW, LB, NR, PD, P-AD, QK, VC, VGri, YY, FWP, VGre; Proteomic analysis, SAA, JK; 3D reconstructions and analysis: ACK, FA, NR, QK, TS, VGri; Electron microscopy, FWP, VD; Ex vivo patch-clamp electrophysiology, AB, ECC, VGre; Ex vivo calcium imaging, AB, CD; In vivo calcium imaging, AA, K-YW, YY; Western blot: LB, M-AC-dS, TL; Behavior analysis: Ace, MK, VGre; Corticosterone dosage, YG, VA, VH; Writing, AB, AC, CPS, FA, FWP, NR, VGre; Project administration and supervision, AC.

