## Extended Data for "Oxytocin Gα_i_ signaling-induced amygdala astrocytes processes retraction shapes behavioral stress response"

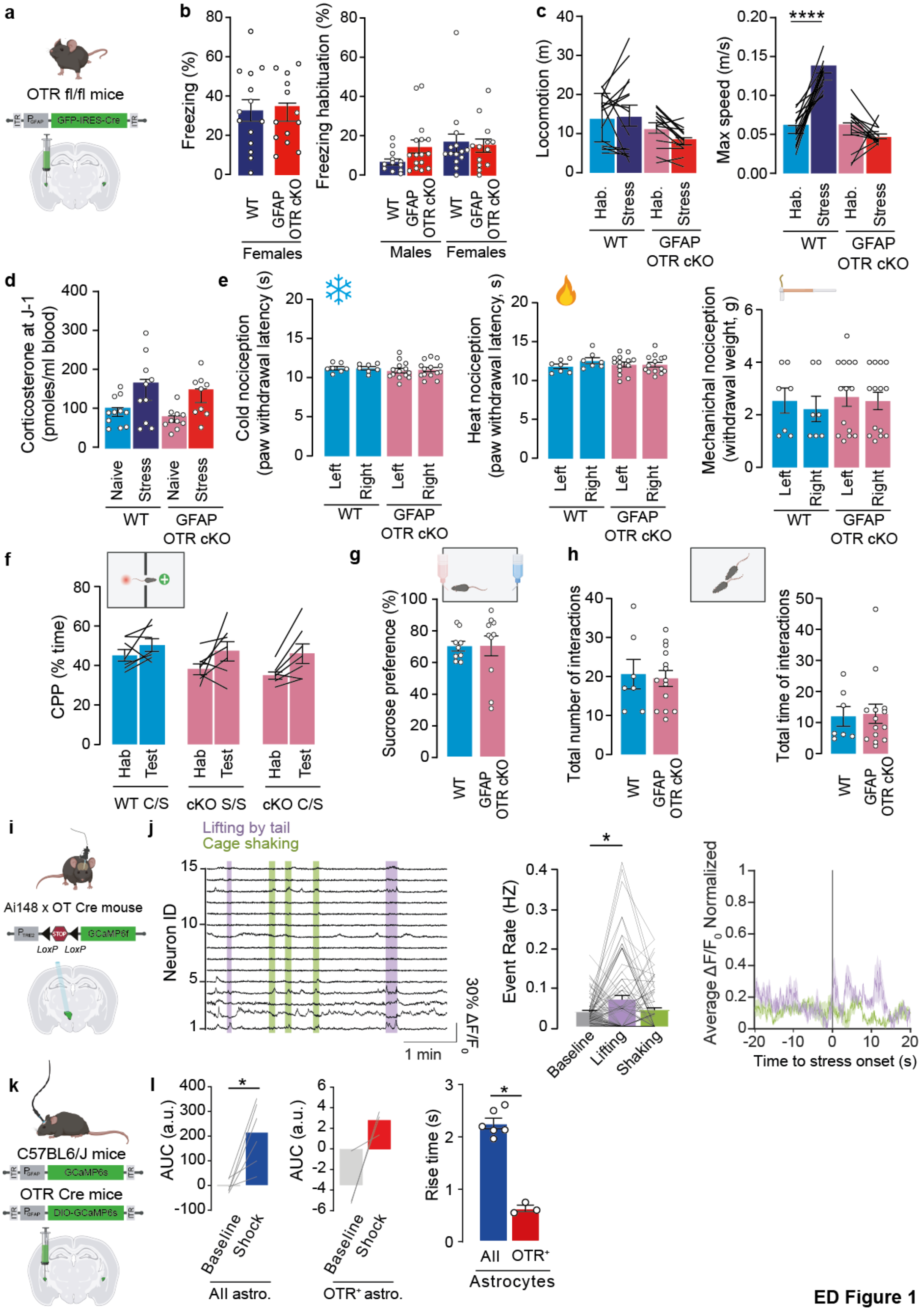
a
OTR fl/fl mice
GFP-IRES-Cre
b
Freezing (%)
WT
GFAP OTR cKO
Females
Freezing habituation (%)
Males
c
Locomotion (m)
Hab.
Stress
Max speed (m/s)
****
d
Corticosterone at J-1 (pmoles/ml blood)
Naive
e
Cold nociception (paw withdrawal latency (s)
Left
Right
Heat nociception (paw withdrawal latency, s)
Mechanichal nociception (withdrawal weight, g)
f
CPP (% time)
Hab
Test
WT C/S
cKO S/S
cKO C/S
g
Sucrose preference (%)
h
Total number of interactions
Total time of interactions
i
Ai148 x OT Cre mouse
GCaMP6f
LoxP
j
Lifting by tail
Cage shaking
Neuron ID
1 min
Event Rate (HZ)
Baseline
Lifting
Shaking
Time to stress onset (s)
k
C57BL6/J mice
GCaMP6s
OTR Cre mice
DIO-GCaMP6s
l
AUC (a.u.)
Shock
All astro.
OTR+ astro.
Rise time (s)
All
OTR+
Astrocytes

**ED Figure 1**

**Extended Data Figure 1: Impact of OTR deletion from CeL/C astrocyte on freezing behavior.**

**a.** GFAP OTR cKO model. AAV-GFAP-Cre was injected in OTR-lox mice to knock out the OTR in astrocytes 3 weeks before stress experiments. **b.** Left: time spent freezing for WT and GFAP OTR cKO female mice. Right: time spent freezing during male and female mice habituation of the stress cage. **c.** Distance travelled (left), and maximum speed (right) before and during the stress protocol. $n_{WT\ males}$=16, $n_{GFAP\ OTR\ cKO\ males}$=13, $n_{WT\ females}$=14, $n_{GFAP\ OTR\ cKO\ females}$=13 mice. **d.** Blood corticosterone levels the day before the stress protocol. $n_{WT\ naïve}$=11, $n_{WT\ stress}$=11, $n_{GFAP\ OTR\ cKO\ naïve}$=10, $n_{GFAP\ OTR\ cKO\ stress}$=10 mice. **e.** Nociceptive characterization of GFAP OTR cKO mice. Left to right: mice response to cold, hot and mechanical stimulation. $n_{WT}$=7, $n_{GFAP\ OTR\ cKO}$=14 mice. **f.** Spontaneous pain assessment. Mice were injected with clonidine, an $\alpha_{2A}$-adrenergic agonist with potent analgesic effect, in a chamber and the conditioned place preference were estimated as the time spent in this chamber. More time spent in the clonidine-injected compartment indicates a clonidine-induced pain relief, thus the presence of spontaneous pain in these animals. In our hand, no spontaneous pain was detected in WT and OTR GFAP cKO. n=7 mice per group. **g.** Anhedonia assessment with sucrose preference test. $n_{WT}$=9, $n_{GFAP\ OTR\ cKO}$=11 mice. **h.** Free social interactions quantification. $n_{WT}$=7, $n_{GFAP\ OTR\ cKO}$=14 mice. **i.** PVN OT neurons calcium activity during different stress. GCaMP6f was expressed in PVN OT neurons by crossbred Ai148 mice with OXT-Cre mice. GRIN lens were implanted above the PVN and GCaMP fluorescence were captured thanks to a miniscope. **j.** Left: example of fluorescence variations in function of time. Each line represent a single neuron. Green vertical lines show cage shaking and purple one indicates when mice were lifted by the tail. Middle: quantification of calcium event rate during different stressors exposure. Right: peristimulus graph showing the mean variation of GCaMP fluorescence in PVN OT neurons before and after different stressors exposure. n=4 mice. **k.** CeL/C astrocyte calcium response temporal features. GCaMP was expressed in all CeA astrocytes by injecting rAAV-GFAP-GCaMP in WT mice (n=5) or only in OTR-expressing CeA astrocytes by injecting rAAV-GFAP-DIO-GCaMP in OTR-Cre mice (n=3), respectively. **l.** Left: quantification of the area under the curve of the fluorescence signal between the baseline of the recording and just after the electric foot shock. Right: rise time of the calcium peak elicited by the electric foot shock. Data are expressed as mean across animal ± SEM. Detailed statistics can be found in *Statistic Table 1*. * p<0.05, ** p<0.01, *** p<0,001, **** P<0.0001.

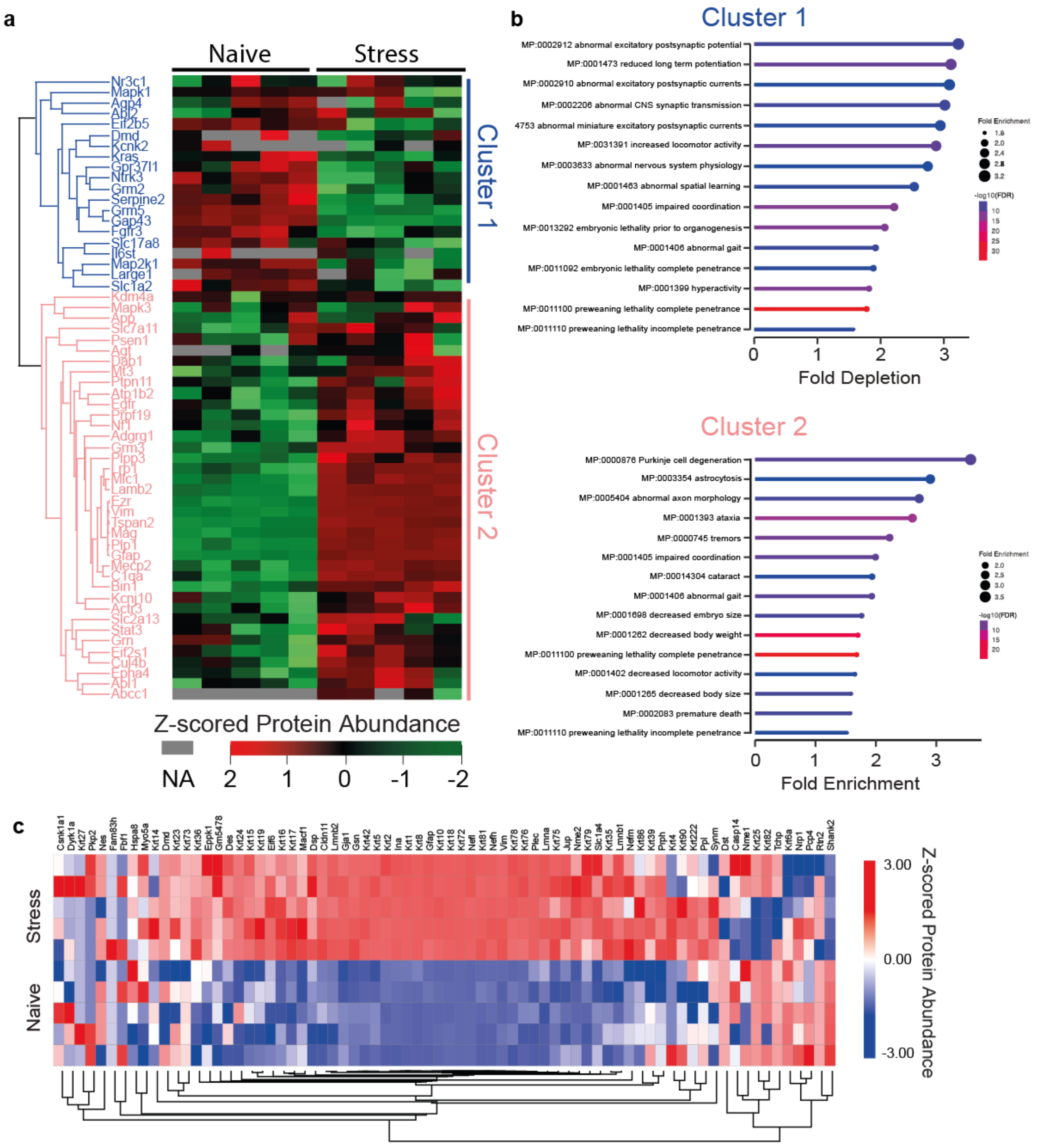

a
Naive
Stress
Nr3c1
Mapk1
Aqp4
Abl2
Eif2b5
Dmd
Kcnk2
Kras
Gpr37l1
Ntrk3
Grm2
Serpine2
Grm5
Gap43
Fgfr3
Slc17a8
Il6st
Map2k1
Large1
Slc1a2
Kdm4a
Mapk3
App
Slc7a11
Psen1
Agt
Dab1
Mt3
Ptpn11
Atp1b2
Egfr
Prpf19
Nf1
Adgrg1
Grm3
Plpp3
Lrp1
Mlc1
Lamb2
Ezr
Vim
Tspan2
Mag
Plp1
Gfap
Mecp2
C1qa
Bin1
Kcnj10
Actb
Slc2a13
Stat3
Grn
Eif2s1
Cul4b
Epha4
Abl1
Abcc1
Cluster 1
Cluster 2
Z-scored Protein Abundance
NA
2
1
0
-1
-2
b
Cluster 1
MP:0002912 abnormal excitatory postsynaptic potential
MP:0001473 reduced long term potentiation
MP:0002910 abnormal excitatory postsynaptic currents
MP:0002206 abnormal CNS synaptic transmission
4753 abnormal miniature excitatory postsynaptic currents
MP:0031391 increased locomotor activity
MP:0003633 abnormal nervous system physiology
MP:0001463 abnormal spatial learning
MP:0001405 impaired coordination
MP:0013292 embryonic lethality prior to organogenesis
MP:0001406 abnormal gait
MP:0011092 embryonic lethality complete penetrance
MP:0001399 hyperactivity
MP:0011100 preweaning lethality complete penetrance
MP:0011110 preweaning lethality incomplete penetrance
0
1
2
3
Fold Depletion
Fold Enrichment
1.6
2.0
2.4
2.8
3.2
-log10(FDR)
10
15
20
25
30
Cluster 2
MP:0000876 Purkinje cell degeneration
MP:0003354 astrocytosis
MP:0005404 abnormal axon morphology
MP:0001393 ataxia
MP:0000745 tremors
MP:0001405 impaired coordination
MP:00014304 cataract
MP:0001406 abnormal gait
MP:0001698 decreased embryo size
MP:0001262 decreased body weight
MP:0011100 preweaning lethality complete penetrance
MP:0001402 decreased locomotor activity
MP:0001265 decreased body size
MP:0002083 premature death
MP:0011110 preweaning lethality incomplete penetrance
0
1
2
3
Fold Enrichment
Fold Enrichment
2.0
2.5
3.0
3.5
-log10(FDR)
10
15
20
c
Stress
Naive
3.00
0.00
-3.00
Z-scored Protein Abundance


**ED Figure 2**

**Extended Data Figure 2: Proteomics analysis of amygdala proteome in control and stressed mice.**

**a.** Heat map and profile plot of astrocyte proteins. Hierarchical clustering was performed for z-scored expression data of astrocyte proteins in the proteomic data of stressed and control mice. Data are displayed as heat map (left panel) and profile plot (right panel). **b.** Top-15 terms obtained by over-representation analysis of Clusters 1 and 2 obtained in Figure 2C. X-axis and circle size indicate fold-enrichment of the term, and color code indicates false discovery rate (FDR). **c.** Heat map of z-scored protein abundance for key cytoskeletal proteins across control and stress conditions, identified by global proteome analysis. Detailed statistics can be found in *Statistic Table 2*.

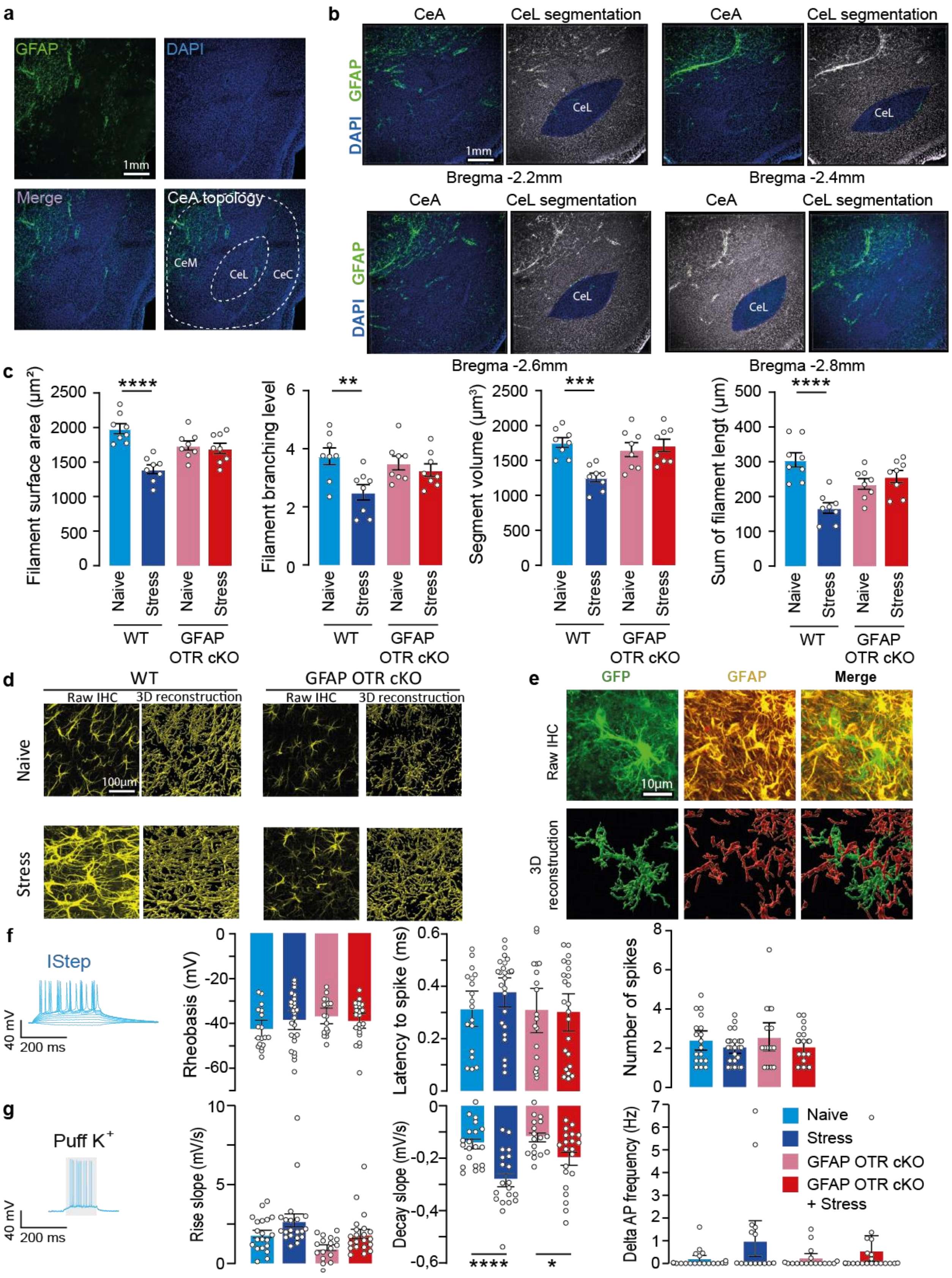
a
GFAP
DAPI
1mm
Merge
CeA topology
CeM
CeL
CeC
b
CeA
CeL segmentation
DAPI GFAP
CeL
1mm
Bregma -2.2mm
Bregma -2.4mm
Bregma -2.6mm
Bregma -2.8mm
c
Filament surface area (µm²)
Filament branching level
Segment volume (µm³)
Sum of filament lengt (µm)
****
**
***
****
Naive
Stress
WT
GFAP OTR cKO
d
WT
GFAP OTR cKO
Raw IHC
3D reconstruction
Naive
Stress
100µm
e
GFP
GFAP
Merge
Raw IHC
3D reconstruction
10µm
f
IStep
40 mV
200 ms
Rheobasis (mV)
Latency to spike (ms)
Number of spikes
g
Puff K+
Rise slope (mV/s)
Decay slope (mV/s)
Delta AP frequency (Hz)
****
*
Naive
Stress
GFAP OTR cKO
GFAP OTR cKO + Stress

ED Figure 3

**Extended Data Figure 3: Morphological analysis of the CeL/C astrocytes network after stress.**

**a.** Overview of mice CeA stained with DAPI (blue) and GFAP (green). Scale bar=1mm. **b.** Segmentation of CeA sub nuclei at different anteroposterior coordinates. Scale bar=1mm. **c.** Left to right: area, filament branching level, volume, and length of GFP-reconstructed astrocytes processes. n=8 mice per group. **d.** Representative low magnification images of raw immunohistochemistry data and 3D reconstruction of astrocytes GFAP labelling in WT and GFAP OTR cKO mice under naïve and stress conditions. Scale bar = 100µm. **e.** Representative images of raw immunohistochemistry data (upper panel) and 3D reconstruction of astrocytes morphology and GFAP labelling (lower panel). Scale bar=10µm. **f.** Features recorded from CeL/C neurons of naïve or stressed WT or GFAP OTR cKO mice during a current step protocol. Left to right: spiking threshold of CeL/C neurons, latency between the beginning of the current step and the first spike, number of spike during the second current step evoking action potentials. **g.** Left to right: rise and decay constant of neuronal membrane depolarization evoked by the puff of a 30mM-containing $K^+$ aCSF, difference of the number of spike before and during the $K^+$ puff. n = 18-21 neurons. Data are expressed as mean across animal ± SEM. Detailed statistics can be found in *Statistic Table 3*. * $p<0.05$, ** $p<0.01$, *** $p<0.001$.

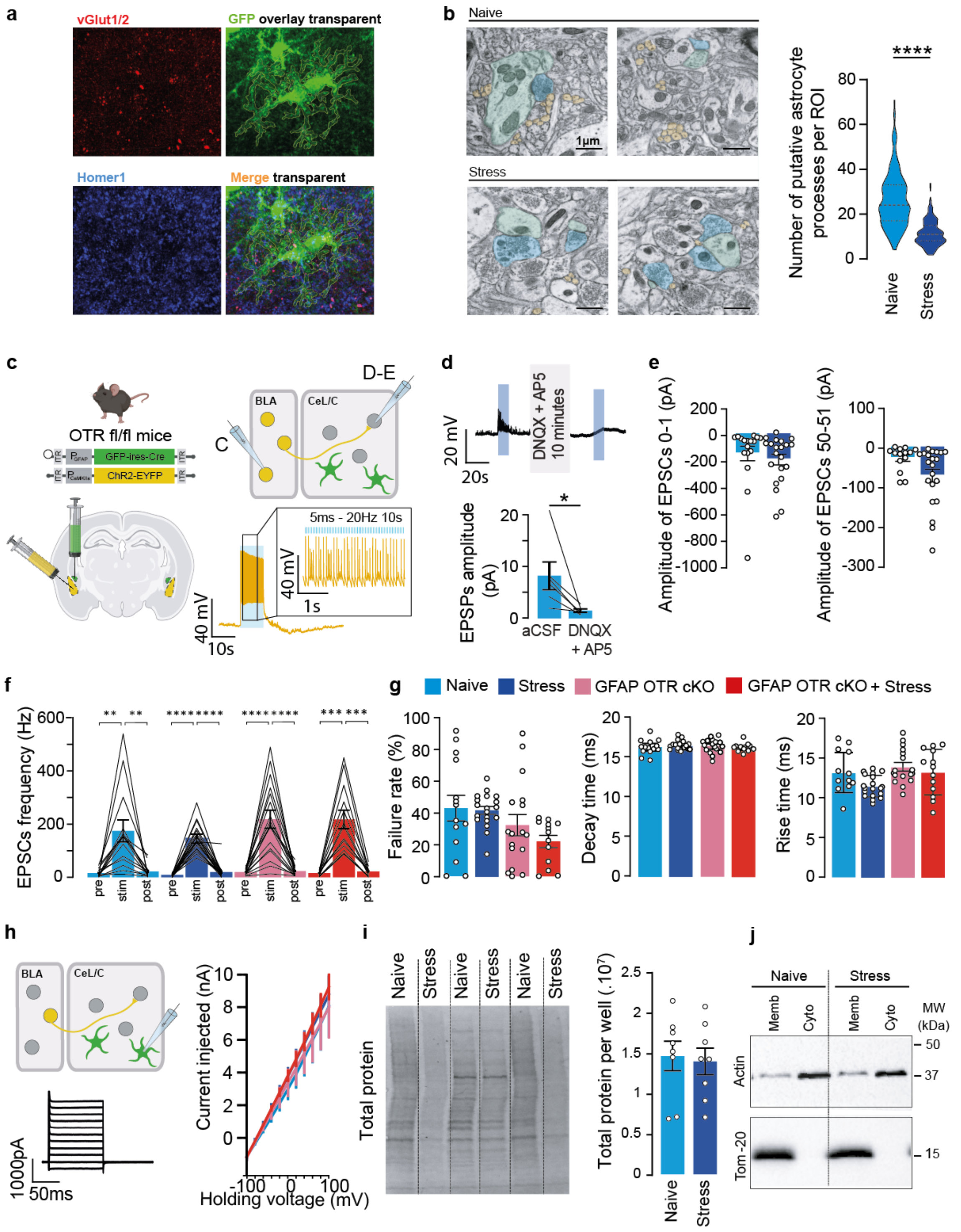

a
vGlut1/2
GFP overlay transparent
Homer1
Merge transparent
b
Naive
Stress
1µm
Number of putative astrocyte processes per ROI
****
Naive
Stress
c
OTR fl/fl mice
GFP-ires-Cre
ChR2-EYFP
D-E
BLA
CeL/C
C
5ms - 20Hz 10s
40 mV
1s
40 mV
10s
d
DNQX + AP5 10 minutes
20 mV
20s
EPSPs amplitude (pA)
*
aCSF
DNQX + AP5
e
Amplitude of EPSCs 0-1 (pA)
Amplitude of EPSCs 50-51 (pA)
f
EPSCs frequency (Hz)
pre
stim
post
g
Naive
Stress
GFAP OTR cKO
GFAP OTR cKO + Stress
Failure rate (%)
Decay time (ms)
Rise time (ms)
h
BLA
CeL/C
1000pA
50ms
Current injected (nA)
Holding voltage (mV)
i
Naive
Stress
Total protein
Total protein per well (.10⁷)
j
Naive
Stress
Memb
Cyto
MW (kDa)
Actin
Tom-20
50
37
15


**ED Figure 4**

**Extended Data Figure 4: Involvement of astrocytic OTR on stress-induced modification of BLA-to-CeL/C excitatory synaptic transmission.**

**a.** Representative fluorescent confocal microscopy images of vGlut1/2 (red) and Homer1 (blue) immunohistochemistery labelling, and GFP (green) fluorescence, as 3D reconstructed in Figure 4. **b.** Left: Representative electron transmission microscopy images depicting a multitude of putative astrocytic processes (yellow) in the vicinity of pre- and post-synaptic elements (respectively blue and green) in naïve but not stressed WT mice. Scale bar=1µm. Right: Quantification of the number of putative astrocytic processes in naïve and stressed WT mice. **c.** Optogenetic control of BLA pyramidal neurons activity. AAV-CaMKII-ChR2 was injected in the BLA of mice. 3 weeks later, we used whole-cell patch-clamp recording in slices to control the ChR2-induced activation of neurons. The yellow trace shows the variation of a BLA neuron membrane potential triggered by blue light pulses (10s of 5ms pulses applied at 20Hz). **d.** To control the glutamatergic nature of the postsynaptic potentials (EPSPs) recorded in CeL/C neurons during ChR2-induced activation of BLA neurons, we injected AAV-CaMKII-ChR2 in the BLA of mice and performed whole-cell patch-clamp recording of CeL/C neurons in slice 3 weeks later. AP5 and DNQX were added in the bath solution for 10 minutes between two photostimulation. Upper panel shows a representative before and after addition of AP5 and DNQX. Lower panel shows a quantification of the effect of the glutamate antagonists AP5 and DNQX on EPSPs amplitudes triggered by ChR2-induced activation of BLA neurons. n=6 neurons. **e.** Quantifications of the BLA ChR2-induced EPSC current amplitudes in CeL/C neurons. **f.** Excitatory postsynaptic currents (EPSCs) frequency recorded in CeL/C neurons before, during and after BLA neurons photostimulation. **g.** Characterization of BLA-to-CeL/C EPSCs triggered by BLA neurons photostimulation. Left to right: failure rate, decay and rise time of CeL/C EPSCs evoked by BLA neurons photostimulation. $n_{WT\ naïve}$=14, $n_{WT\ stress}$=19, $n_{GFAP\ OTR\ KO\ naïve}$=17, $n_{GFAP\ OTR\ KO\ stress}$=13 neurons. **h.** To control the astrocytic nature of recorded cells, we performed a voltage-current relationship recorded in astrocytes from naïve and stressed mice. $n_{WT\ naïve}$=13, $n_{WT\ stress}$=11, $n_{GFAP\ OTR\ KO\ naïve}$=11, $n_{GFAP\ OTR\ KO\ stress}$=10 astrocytes. **i.** Left: total protein load per well was detected via UV light on Stain Free gels in CeA samples from stress and naïve mice. Right: quantification of the total protein content in stress and naïve samples. These values were used for normalization in Figure 5, as per using Stain Free gels. n=8 in each condition. **j.** Enrichment in membrane versus cytosolic fractions was confirmed by detecting actin (enriched the cytosolic fraction) and Tom-20 (a mitochondrial membrane protein, enriched in the membrane fraction). Data are expressed as mean across cells ± SEM for panel A to F, and as mean across animal ± SEM for panels G to J. Detailed statistics can be found in *Statistic Table 5*. * p<0.05, ** p<0.01.

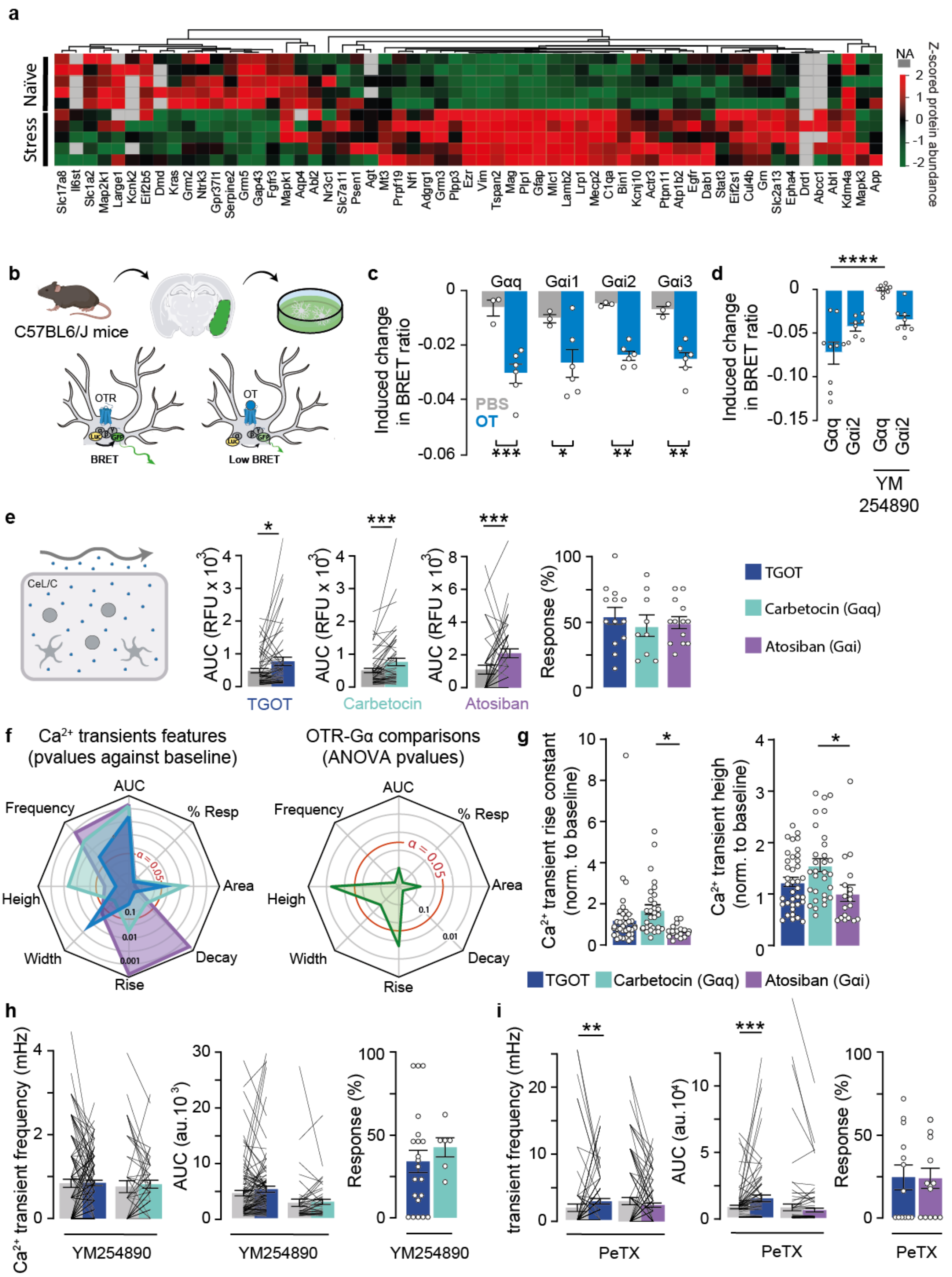
a
Naïve
Stress
NA
Z-scored protein abundance
Slc17a8
Il6st
Slc1a2
Map2k1
Large1
Kcnk2
Eif2b5
Dmd
Kras
Grm2
Ntrk3
Gpr37l1
Serpine2
Grm5
Gap43
Fgfr3
Mapk1
Aqp4
Abl2
Nr3c1
Slc7a11
Psen1
Agt
Mt3
Prpf19
Nf1
Adgrg1
Grm3
Plpp3
Ezr
Vim
Tspan2
Mag
Plp1
Gfap
Mlc1
Lamb2
Lrp1
Mecp2
C1qa
Bin1
Kcnj10
Actr3
Ptpn11
Atp1b2
Egfr
Dab1
Stat3
Eif2s1
Cul4b
Gm
Slc2a13
Epha4
Drd1
Abcc1
Abl1
Kdm4a
Mapk3
App
b
C57BL6/J mice
OTR
OT
BRET
Low BRET
c
Gαq
Gαi1
Gαi2
Gαi3
Induced change in BRET ratio
PBS
OT
d
YM 254890
e
CeL/C
AUC (RFU x 10^3)
Response (%)
TGOT
Carbetocin
Atosiban
TGOT
Carbetocin (Gαq)
Atosiban (Gαi)
f
Ca2+ transients features (pvalues against baseline)
OTR-Gα comparisons (ANOVA pvalues)
AUC
% Resp
Area
Decay
Rise
Width
Heigh
Frequency
α = 0.05
g
Ca2+ transient rise constant (norm. to baseline)
Ca2+ transient heigh (norm. to baseline)
TGOT
Carbetocin (Gαq)
Atosiban (Gαi)
h
Ca2+ transient frequency (mHz)
AUC (au.10^3)
Response (%)
YM254890
i
transient frequency (mHz)
AUC (au.10^4)
Response (%)
PeTX

ED Figure 5

**Extended Data Figure 5: Intracellular pathway activated downstream OTR.**

**a.** Differential expression of cAMP-related signaling proteins in the amygdala under control and stress conditions. Heat map of Z-scored protein abundance of cAMP-related signaling proteins identified in the amygdala proteome from naïve and stressed mice. **b.** Verification of the biased OTR activation triggered by atosiban and carbetocin. Right: primary astrocyte cultures were obtained from temporal lobe of mice brains. Left: BRET donors and acceptors were expressed in cultured astrocytes. The BRET donor Rluc8 was inserted in the indicated different Gα proteins and the BRET acceptor GFP[10] was attached to the $G\gamma_2$ protein, allowing to monitor their activation, measured by a decrease of BRET ratio. **c.** Induced changes in BRET ratio due to OTR activation by OT in live astrocytes. $n_{PBS}$=3 and $n_{OT}$=6 in each condition. **d.** Verification of YM254890 ability to prevent OTR/Gαq without changing OTR/Gαi2 interaction measured as BRET ratio in HEK cells. $n_{G\alpha q}$=9, $n_{G\alpha i2}$=8. **e.** Calcium imaging of SR101-identified CeL/C astrocytes somata before and after bath application of the biased agonists. Left: area under the curve of calcium imaging traces. Right: proportion of responsive astrocytes per recording to agonist application. $n_{TGOT}$=57 astrocytes over 13 slices, $n_{atosiban}$=49 astrocytes over 9 slices, $n_{carbetocin}$=54 astrocytes over 13 slices. **f.** Comparison of calcium activity triggered by different OTR agonists. Spider plots showing (left) the p-value of the comparison between the basal state and after the application of different agonists for different calcium signaling parameters and (right) the p-value of a one-way ANOVA to compare calcium events triggered by the three different agonists. **g.** Bar plots showing mean rise time and height of calcium transients triggered by indicated OTR agonists normalized to baseline values **h.** Frequency and area under the curve of calcium transients, and proportion of responsive cells to carbetocin and TGOT after acute brain slices incubation in YM254890 to block Gαq-dependent signaling. $n_{TGOT}$=110 astrocytes over 21 slices, $n_{carbetocin}$=48 astrocytes over 6 slices. **i.** Frequency and area under the curve of calcium transients, and proportion of responsive cells to different atosiban and TGOT after acute brain slices incubation in pertussis toxin (PeTX) to block Gαi -dependent signaling. $n_{TGOT}$=90 astrocytes over 13 slices, $n_{atosiban}$=99 astrocytes over 14 slices. Data are expressed as mean across cell ± SEM. Detailed statistics can be found in *Statistic Table 6*. * $p<0.05$, ** $p<0.01$, *** $p<0.001$.

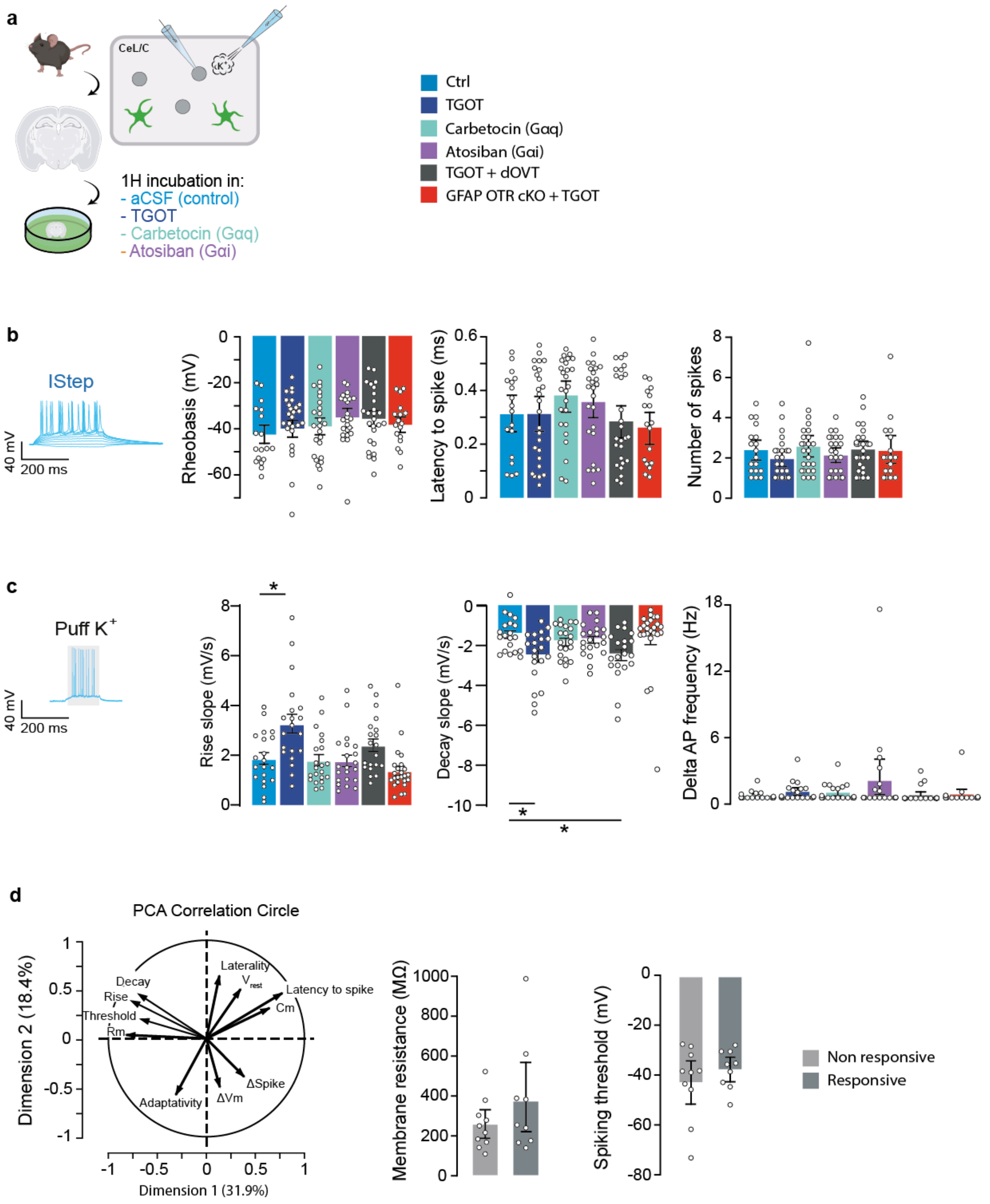

a
CeL/C
1H incubation in:
- aCSF (control)
- TGOT
- Carbetocin (Gαq)
- Atosiban (Gαi)
Ctrl
TGOT
Carbetocin (Gαq)
Atosiban (Gαi)
TGOT + dOVT
GFAP OTR cKO + TGOT
b
IStep
40 mV
200 ms
Rheobasis (mV)
Latency to spike (ms)
Number of spikes
c
Puff K+
Rise slope (mV/s)
Decay slope (mV/s)
Delta AP frequency (Hz)
d
PCA Correlation Circle
Dimension 2 (18.4%)
Dimension 1 (31.9%)
Laterality
Decay
Rise
Threshold
Rm
Latency to spike
Cm
ΔSpike
ΔVm
Adaptativity
Membrane resistance (MΩ)
Spiking threshold (mV)
Non responsive
Responsive


**ED Figure 6**

**Extended Data Figure 6: Effect of the different OTR intracellular coupling on the astro-neuronal network.**

**a.** Brain slices were incubated for 1 hour in the different OTR agonists before electrophysiological characterization of CeL/C neurons. **b.** Left to right: spiking threshold of CeL/C neurons, latency between the beginning of the current step and the first spike, number of spike during the second current step evoking action potentials. **c.** Left to right: rise and decay constant of neuronal membrane depolarization evoked by the puff of a 30mM-containing $K^+$ aCSF. Difference of the number of spike before and during the $K^+$ puff. **d.** Upon closer examination of neuronal $K^+$ sensitivity, we observed that certain neurons exhibited heightened membrane depolarization in response to $K^+$ puff, while this reaction remained unchanged in others. To explore the presence of a distinct subset of neurons displaying increased sensitivity to extracellular $K^+$ after OT stimulation, we conducted a principal component analysis on the electrophysiological properties of CeL/C neurons. This analysis revealed that the first principal components explained 50.3% of the variability in the data (PC1: 31.9%, PC2: 18.4%), highlighting a negative correlation between neuronal response to $K^+$ and two passive membrane characteristics: the spiking threshold and membrane resistance of neurons. However, the comparisons of these parameters were not significant, indicating that they cannot predict neuron's hypersensitivity to extracellular $K^+$ following TGOT incubation ($p_{threshold}$=0.351, $p_{Rm}$=0.240). Left: PCA correlation circle showing a positive correlation between depolarisation amplitude triggered by $K^+$ puff the number of spike it elicits. The circle also show an inverse correlation between depolarisation amplitude, its rise & decay constants, the membrane resistance of the neuron and its spiking threshold. Middle & right: Neurons were classified as responsive or non-responsive based on their depolarisation amplitude triggered by $K^+$ puff. The membrane resistance and the spiking threshold were compared between responsive cells and non-responsive cells. n=20 neurons. Data are expressed as mean across neurons ± SEM. Detailed statistics can be found in *Statistic Table 7*. * p<0.05.

### Extended Data Table 1 : List of reagents

| Experiments | Antigen | Host | Company | Company reference | Fluorophore | Concentration final | medium |
|---|---|---|---|---|---|---|---|
| Immunohistochemistry | GFAP (morphology) | Goat | Abcam | ab53554 | / | 1/200 | PBS |
| | GFAP (cFos) | Mouse | Sigma-Aldrich | G3893 | | 1/400 | |
| | cFos | Rabbit | Abcam | ab190289 | / | 1/500 | |
| | S100beta | Mouse | Abcam | ab11178 | / | 1/1000 | |
| | GFP | Chicken | Aves | 1020 | / | 1/200 | |
| | Homer1 | Rabbit | Synaptic system | 160003 | / | 1/200 | |
| | vGlut1 | Mouse | Synaptic system | 135511 | / | 1/200 | |
| | vGlut2 | Mouse | Synaptic system | 135421 | / | 1/200 | |
| | Anti-mouse | Goat | Abberior | STRED-1001-500UG | Star Red | 1/200 | |
| | Anti-mouse | Sheep | Sigma-Aldrich | C2181 | Cyanin3 546 | 1/500 | |
| | Anti-rabbit | Donkey | Interchim | 715-586-152 | Alexa 594 | 1/200 | |
| | Anti-rabbit | Goat | Dianova | 111-065-003 | Biotin-SP | 1/500 | |
| | Avidin anti biotin | | Invitrogen | A21370 | Alexa 488 | 1/1000 | |
| **Experiments** | **Virus name** | **Full name** | **Company** | **Company reference** | **Concentration** | **Injected volume** | **medium** |
| Viruses stereotaxic injections | hSyn-DIO-GCaMP6f | AAV1-hSyn-DIO-GCaMP8m-WPRE | IGBMC | A286 | 1E+13 | 200 nL | PBS |
| | GFAP-GCaMP6s | AAV1.GfaABC1D-GCaMP6s | IGBMC | A238 | 1E+13 | 200 nL | |
| | GFAP-DIO-GCaMP6s | AAV1.GfaABC1D-DIO-GCaMP6s | IGBMC | A247 | 1E+13 | 200 nL | |
| | GFAP-eGFP | pAAV5.GfaABC1D.PI.Lck-GFP.SV40 | Addgene | 105598-AAV5 | 1E+13 | 200 nL | |
| | GFAP-GFP-IRES-Cre | rAAV1/2-gfaABC1D-GFP-IRES-Cre | IGBMC | A270 | 1E+13 | 200 nL | |
| | EF1a-DIO-GFP | AAV5-EF1a-DIO-GFP | IGBMC | A143 | 1E+13 | 200 nL | |
| | CaMKIIa-ChR2-EYFP | AAV9-CaMKIIa-hChR2(h134R)_EYFP | Addgene | 26969-AAV9 | 1E+13 | 100 nL | |
| **Experiments** | **Name** | **Full name** | **Company** | **Company reference** | **Concentration** | **medium** | |
| | HCl | Hydrochloric acid | Termo fischer scientific | H1200PB15 | 93 | aCSF | |
| | NMDG | N-methyl-D-glucamine | Sigma-Aldrich | M2004 | 93 | | |

| | | | | | | |
|---|---|---|---|---|---|---|
| | KCl | Potassium chloride | Carlo erba | 471177 | 2,5 | |
| | NaH2PO4 | Sodium phosphate monobasic monohydrate | Sigma-Aldrich | S9638 | 1,25 | |
| | NaHCO3 | Sodium bicarbonate | Sigma-Aldrich | S5761 | 30 | |
| | HEPES | 4-(2-Hydroxyethyl)piperazine-1-ethanesulfonic acid, N-(2-Hydroxyethyl)piperazine-N′-(2-ethanesulfonic acid) | Sigma-Aldrich | H4034 | 20 | |
| | D-Glucose | Dextrose Glucose | Sigma-Aldrich | G8270 | 25 | |
| | L-Ascorbic Acid | L-Threoascorbic acid | Sigma-Aldrich | A92902 | 5 | |
| | Thiourea | | Sigma-Aldrich | T8656 | 2 | |
| | Sodium Pyruvate | ketopropionic acid sodium salt | Sigma-Aldrich | P2256 | 3 | |
| | N-Acetyl-L-Cysteine | | Sigma-Aldrich | A7250 | 10 | |
| | Kynurenic acid | | Sigma-Aldrich | K3375 | 2,5 | |
| | MgSO4.7H2O | Magnesium sulfate hydrate | Merck | 105886 | 10 | |
| | CaCl2.2H20 | | | | 0,5 | |
| **Calcium imaging on slices** | Kolliphor® EL | Polyoxyl 35 hydrogenated castor oil | Sigma-Aldrich | C5135 | | aCSF Bath medium |
| | OGB1-AM | Oregon Green BAPTA 1 acetoxymethylesther | Thermo Fisher Scientific | O6807 | | |
| | Rhodamine 2 | / | | R1245MP | | |
| | Pluronic F127 | / | Sigma-Aldrich | P2443 | | |
| | Sulforhodamine 101 | / | | S7635 | 1 µM in ACSF | |
| **Patch-Clamp recordings** | Various salts and other products for intracellular solution | / | Sigma-Aldrich | see company website | see materials and methods | |
| | BAPTA | 1,2-Bis(2-Aminophenoxy)ethane-N,N,N′,N′-tetraacetic acid | | A4926 | see materials and methods | |
| | dOVT | (d(CH$_2$)$_5^1$,Tyr(Me)$^2$,Thr$^4$,Orn$^8$,des-Gly-NH$_2^9$)-Vasotocin trifluoroacetate salt | Bachem | H2908 | 1 µM in ACSF | |
| | TGOT | (Thr$^4$,Gly$^7$)-Oxytocin ; H-Cys-Tyr-Ile-Thr-Asn-Cys-Gly-Leu-Gly-NH$_2$ | Bachem | H-7710 | 0,4 µM in ACSF | |
| | TTX citrate | Octahydro-12-(hydroxymethyl)-2-imino-5,9:7,10a-dimethano-10aH-[1,3]dioxocino[6,5-d] pyrimidine-4,7,10,11,12-pentol + citrate buffer | Abcam | ab120055 | 1 µM in ACSF | |
| | K-gluconate | 2,3,4,5,6-Pentahydroxycaproic acid potassium salt | Sigma-Aldrich | G4500 | 125 | Intrapipette solution |

| | | | | |
|---|---|---|---|---|
| EGTA | Ethylene glycol-bis(2-aminoethylether)-*N,N,N′,N′*-tetraacetic acid | Sigma-Aldrich | E3889 | 5 |
| HEPES | 4-(2-Hydroxyethyl)piperazine-1-ethanesulfonic acid, N-(2-Hydroxyethyl)piperazine-N′-(2-ethanesulfonic acid) | Sigma-Aldrich | H4034 | 20 |
| MgCl2 | Magnesium chloride | Sigma-Aldrich | M8266 | 1,3 |
| NaCl | Sodium Chloride | Sigma-Aldrich | 793566 | 10 |
| GTPNa3 | Guanosine 5′-triphosphate sodium salt hydrate | Sigma-Aldrich | G8877 | 0,4 |
| ATPNa2 | Adenosine 5′-triphosphate disodium salt hydrate | Sigma-Aldrich | A2383 | 4 |

### Extended Data Table 2: HPLC and MSMS details

**Elution gradient for corticosterone assessment through HPLC column.**

| **Time (min)** | 0 | 1 | 3 | 10 | 12 | 14 | 15 | 19 |
|---|---|---|---|---|---|---|---|---|
| **% B mobile phase** | 0 | 0 | 25 | 30 | 98 | 98 | 0 | 0 |

**Detail of mass spectrometry analysis conditions.**

| | |
|---|---|
| **Mode** | Positive |
| **Spray voltage** | 3,500 V |
| **Nebulizer gas** | Nitrogen |
| **Desolvation (nitrogen) sheath gas** | 20 Arb |
| **Aux gas** | 7 Arb |
| **Vaporizer Temp (°C)** | 144 |
| **Ion transfer tube temperature** | 299°C. |
| **Q1 and Q2 resolutions** | 0.7 FWHM, |
| **Collision gas (CID, argon) pressure** | 2 mTorr |

**Mass spectrometer ionization, selection, fragmentation, and identification parameters.**

| **Compound** | **Polarity** | **Precursor (m/z)** | **Product (m/z)** | **Collision Energy (V)** | **RF Lens (V)** |
|---|---|---|---|---|---|
| **Corticosterone** | positive | 347,11 | 293,472<br>311,294<br>329,169 | 17,03<br>15,91<br>14,95 | 227,49 |
| **D4-corticosterone** | positive | 351,179 | 297,103<br>315,183<br>333,24 | 17,68<br>16,88<br>15,56 | 223,55 |
